# Evolution of gene expression across brain regions in behaviorally divergent deer mice

**DOI:** 10.1101/2023.09.28.559844

**Authors:** Andreas F. Kautt, Jenny Chen, Caitlin L. Lewarch, Caroline Hu, Kyle Turner, Jean-Marc Lassance, Felix Baier, Nicole L. Bedford, Andres Bendesky, Hopi E. Hoekstra

## Abstract

The evolution of innate behaviors is ultimately due to genetic variation likely acting in the nervous system. Gene regulation may be particularly important because it can evolve in a modular brain-region specific fashion through the concerted action of *cis*- and *trans*-regulatory changes. Here, to investigate transcriptional variation and its regulatory basis across the brain, we perform RNA sequencing (RNA-Seq) on ten brain subregions in two sister species of deer mice (*Peromyscus maniculatus* and *P. polionotus*) – which differ in a range of innate behaviors, including their social system – and their F_1_ hybrids. We find that most of the variation in gene expression distinguishes subregions, followed by species. Interspecific differential expression (DE) is pervasive (52–59% of expressed genes), whereas the number of DE genes between sexes is modest overall (∼3%). Interestingly, the identity of DE genes varies considerably across brain regions. Much of this modularity is due to *cis*-regulatory divergence, and while 43% of genes were consistently assigned to the same gene regulatory class across subregions (e.g., conserved, *cis*-, or *trans*-regulatory divergence), a similar number were assigned to two or more different gene regulatory classes. Together, these results highlight the modularity of gene expression differences and divergence in the brain, which may be key to explain how the evolution of brain gene expression can contribute to the astonishing diversity of animal behaviors.

## Introduction

Differences in innate behaviors are observed even among closely-related species, from feeding and reproductive behavior in *Drosophila* (Markow and O’Grady 2008), to sleep behavior in cavefish (Duboué, Keene, and Borowsky 2011), to social behaviors in mammals (Young et al. 1999). Changes that affect gene expression in the nervous system are thought to be particularly relevant in the evolution of behavioral divergence (Alaux et al. 2009; Barrett et al. 2013; Pizzollo, Zintel, and Babbitt 2022). Thus, profiling gene expression changes across species can be an important first step in elucidating the molecular mechanisms underlying the evolution of behaviors.

Deer mice of the genus *Peromyscus* occupy almost every terrestrial ecological niche in North America, likely contributing to the large amount of interspecific variation in naturally-occurring behaviors (Bedford and Hoekstra 2015; Dewey and Dawson 2001). For example, two sister species, *P. maniculatus* and *P. polionotus*, diverged from each other around 1.8 million years ago (Schenk, Rowe, and Steppan 2013) (Fig. 1A) but nonetheless vary in a number of innate behaviors (e.g., burrowing (Dawson, Lake, and Schumpert 1988), infant vocalization (Jourjine et al. 2023), and thermoregulatory nesting (Lewarch and Hoekstra 2018)). In particular, these two species lie on opposite ends of the monogamy-promiscuity spectrum and, among other reproductive behaviors, exhibit differences in the degree of sexual dimorphism in parental care (Bendesky et al. 2017; Foltz 1981). Because these two species can be interbred to produce F_1_ hybrids, this system provides an opportunity to not only compare interspecific differences in gene expression but also to better understand the regulatory mechanisms underlying those differences.

**Fig. 1.**
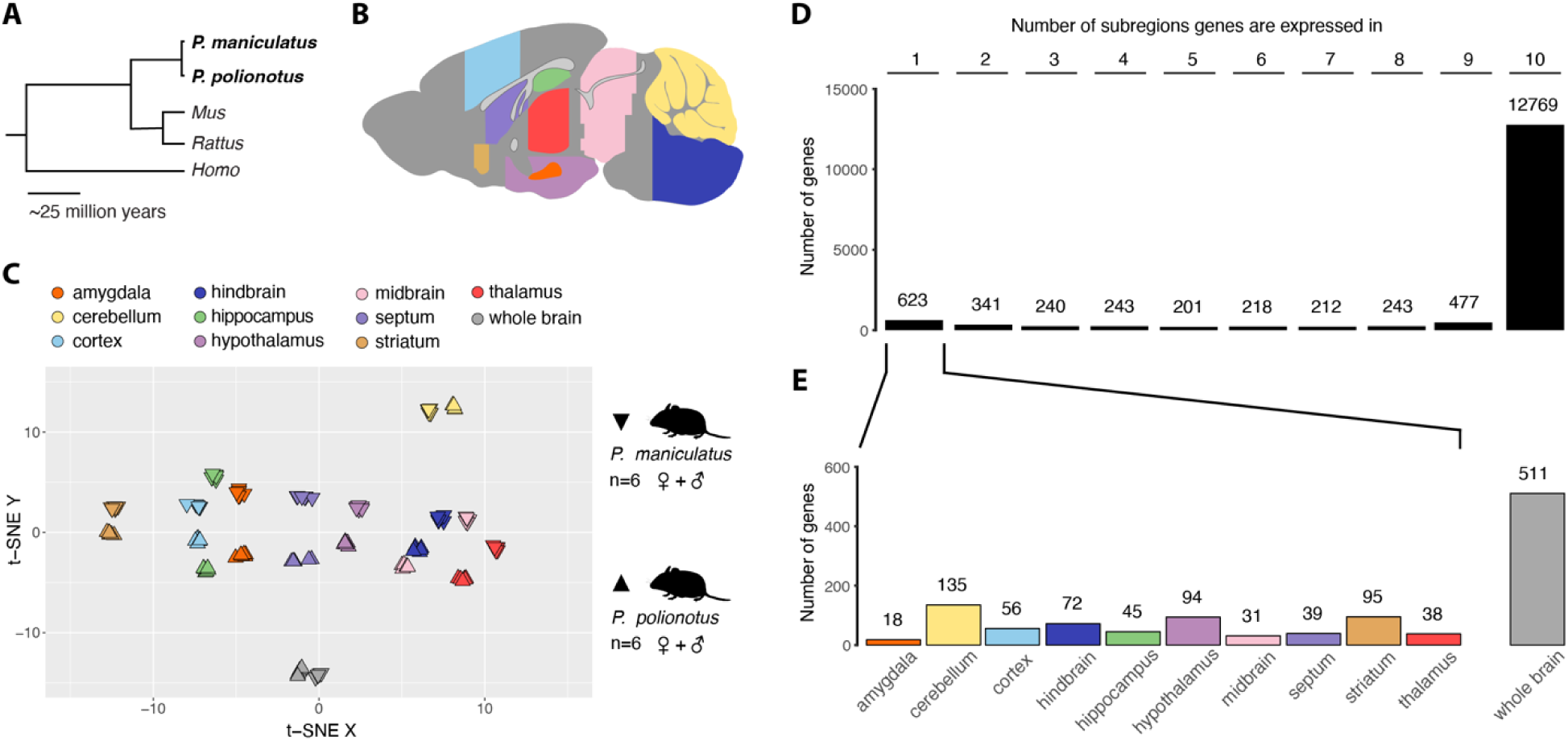
Gene expression across the brain of *P. maniculatus* and *P. polionotus* mice. (**A**) Phylogenetic relationships of the two focal *Peromyscus* species (in bold) with house mice (*Mus*), rats *(Rattus*), and humans (*Homo*). (**B**) Schematic of a sagittal brain section highlighting the locations of the 10 (sub)region dissections (color) used in this study, as well as whole brain (grey). For details on dissections, see Materials & Methods and Table S2. (**C**) t-distributed stochastic neighbor embedding (t-SNE) plot of the overall variation in gene expression. *P. maniculatus* samples are depicted as downward-facing triangles, *P. polionotus* samples as upward-facing triangles. (**D**) The number of genes (n=16,078) expressed in one or multiple brain regions. (**E**) Distribution of privately expressed genes across the 10 (sub)regions; 511 genes are expressed in the whole brain samples but not in any of the 10 sampled (sub)regions.

Comparing species expression differences to allelic expression differences in the F_1_ hybrids allows one to dissect the *cis*- (i.e., genetic variants residing on the same chromatid) and *trans*-regulatory (i.e., genetic variants affecting diffusible elements) components of expression divergence because both alleles in F_1_ hybrids share the same *trans*-regulatory environment (Goncalves et al. 2012; McManus et al. 2010; Tirosh et al. 2009; Verta and Jones 2019; Wittkopp, Haerum, and Clark 2008). While a few studies have used F_1_ hybrids to profile regulatory changes in the brains of behaviorally divergent species (Benowitz et al. 2020; Reuveni et al. 2018; Wang et al. 2019), they have typically focused on only one or two brain regions of interest. Given that the brain is a heterogeneous organ composed of functionally specialized subregions composed of transcriptionally distinct cell clusters (Zhang et al. 2023), this raises the question: if a gene is differentially expressed in one region of the brain, is it typically also differentially expressed in other regions, and what is the regulatory basis underling this modularity in gene expression?

To address this question and gain a better understanding of the evolution of gene regulation in the brain, we investigate gene expression in both females and males of *P. maniculatus* and *P. polionotus* as well as their F_1_ hybrids using RNA-Seq. We compare variation in gene expression in the whole brain as well as across ten distinct subregions of the brain: the amygdala, cerebellum, cortex, hindbrain, hippocampus, hypothalamus, midbrain, septum, striatum, and the thalamus (Fig. 1B, Fig. S1). Using these data, we (i) quantify gene expression differences across the brain, (ii) screen for differentially expressed genes between sexes and species, and (iii) investigate the regulatory mechanisms driving this gene expression divergence. Importantly, in addition to detecting and characterizing DE genes *per se*, we also (iv) assess the extent of shared versus region-specific gene expression differences and their underlying regulatory divergence across the brains of these two deer mice species.

## Materials and Methods

### Animal husbandry

*P. maniculatus bairdii* and *P. polionotus subgriseus* animals were originally acquired from the *Peromyscus* Genetic Stock Center (Columbia, SC, USA). To generate F_1_ hybrids, we crossed female *P. maniculatus* and male *P. polionotu*s, since hybrids derived from reciprocal crosses are inviable (Vrana et al. 2000; Watson 1942). We weaned animals at 21–23 days of age into single-sex, same-strain groups. We maintained animals on a 16 h:8 h light:dark cycle at 22°C, housed them in standard mouse cages with corncob bedding and provided them with food and water *ad libitum*. Animal husbandry and experimental procedures were approved by the Harvard University Faculty of Arts and Sciences Institutional Animal Care and Use Committee (protocol 27-15-3).

### Tissue collection, RNA extraction, library preparation, and sequencing

#### Whole brain data

For the whole brain dataset, a cohort of 59–137-day-old virgin male *P. maniculatus* (n=5) and *P. polionotus* (n=6) animals were euthanized by CO_2_ inhalation 7–10 h before the start of the dark period. Brains were then rapidly dissected, flash-frozen, and stored at -70°C. We homogenized the samples in TRIzol Reagent (Invitrogen, 15596026) using a Bio-Gen PRO200 homogenizer (PRO Scientific, Oxford, USA), and extracted RNA using a Direct-zol™ RNA MiniPrep Plus kit (Zymo Research, R2070), including a DNase I treatment. We measured RNA concentrations using a Qubit RNA BR Assay kit (Thermo Fisher Scientific, Q10210) and assessed RIN scores using an Agilent TapeStation 2200 (Agilent Technologies, Santa Clara, CA).

We prepared each sequencing library from 1 μg of total RNA using a PrepX PolyA mRNA Isolation Kit (Wafergen, 40047) on the Apollo 324 NGS Library Prep System (Takara Bio USA Holdings, Mountain View, CA). We added Illumina indices and amplified libraries with 12 cycles of PCR and cleaned them using PCRClean DX beads (Aline Biosciences, Woburn, MA). To quantify library concentrations, we used the Quant-iT BR dsDNA kit (ThermoFisher Scientific, Q33130). We then assessed library fragment size distributions using an Agilent TapeStation 2200 D1000 tape. To further quantify the libraries, we used a Kapa Library Quantification Kit for Illumina Platforms (Kapa Biosystems, Wilmington, MA). In total, we pooled 33 libraries to 20 nM and sequenced them on 5 lanes of an Illumina HiSeq 2500 2×150 bp chemistry. Additional information on samples and sequencing batches is provided in Table S1 and Fig. S2B.

#### Subregion data

For the brain subregion dataset, a cohort of 82–85-day-old virgin male and female *P. maniculatus, P. polionotus,* and their F_1_ hybrids (see Table S1 for sample sizes) were euthanized by CO_2_ inhalation followed by cervical dislocation 3–10 h before the onset of the dark period. One cohort was used for dissection of the cerebellum, hippocampus, and hindbrain samples. In these samples, after the brain was removed, a dorsal-to-ventral incision was made posterior to the cerebrum to release the hindbrain and cerebellum. The cerebellar peduncles were cut, and the hindbrain and cerebellum were placed in tubes and flash frozen in liquid nitrogen. We note that our dissection strategy may have inadvertently captured part of the midbrain in the hindbrain samples. The petri dish with the remaining brain was then placed on ice, and the two cerebral hemispheres were cut apart. For each hemisphere separately, the hippocampus was severed from the dorsal fornix and gently scooped out of the hemisphere with a brush in ventral-to-dorsal direction (Lu, Airey, and Williams 2001), and then flash frozen. For a second cohort of animals, we dissected the brains and then immediately submerged them into embedding compound for cryosectioning (Tissue-Tek O.C.T.) and froze them on dry ice. We sectioned brains on a coronal plane at 150 μm using a cryostat at -20°C (Leica CM3050 S, Leica Biosystems Inc., Buffalo Grove, IL) and stored the sections at -70°C until we dissected the medial prefrontal cortex, septum, striatum, hypothalamus, amygdala, thalamus, and midbrain with sample corers according to anatomical landmarks (Fig. S1, Table S2). The sample cores were immediately transferred into homogenization buffer. After subregion dissection, we homogenized samples for 10 seconds in Maxwell RSC Homogenization Buffer (Promega, Madison, USA) with 2% 1-thioglycerol using a Bio-Gen PRO200 homogenizer (PRO Scientific, Oxford, USA) and then extracted RNA using the Maxwell RSC simplyRNA Tissue Kit (Promega). We quantified extracts using a Quant-IT RNA Assay Kit, assayed for purity on a NanoDrop ND-1000, and tested for RNA integrity on a TapeStation 4200.

We prepared RNA-Seq libraries using an mRNA HyperPrep kit (Kapa Biosystems, Wilmington, USA) automated on a BioMek FXp (Beckman Coulter, Brea, USA). Post-capture, we fragmented mRNA for 6 min at 94°C for a target insert size of 200-300 base pairs. We amplified dual-indexed adapter-ligated libraries in 11 PCR cycles and then sequenced the amplified libraries on a NovaSeq 6000 2×150 bp on two S2 flow cells.

### Trimming and read mapping

Using *cutadapt v.2.3* (Martin 2011), we trimmed very low quality bases (quality score <10) and applied a minimum length cutoff of 25 bp. We then mapped trimmed reads of *P. maniculatus* and *P. polionotus* individuals to their respective reference genome, Pman_2.1 (GCA_003704035.1) or Ppol_1.3.3 (GCA_003704135.2), and in-house annotations generated with Comparative Annotation Toolkit (Fiddes et al. 2018) using the GENCODE v15 primary-assembly annotation of *Mus musculus* (GRCm38 mm10) as reference. We competitively mapped reads from F_1_ hybrids to a diploid reference genome created by concatenating the two species assemblies and annotations. We performed read mapping with *STAR v.2.7.0e* (Dobin et al. 2013) in a transcriptome-aware manner. On average, we successfully mapped 30.18 ± 5.44 (SD), 31.58 ± 6.99 (SD), and 63.06 ± 15.34 (SD) million read pairs per sample for *P. maniculatus, P. polionotus*, and their F_1_ hybrids, respectively (Table S3).

### Gene expression estimation

We estimated gene expression with *RSEM v1.3.1* (Li and Dewey 2011). We retained only genes present in both species’ annotations for downstream analyses (n = 33,836). We performed a blast search against an NCBI-generated annotation of *P. maniculatus* (Annotation Release 100) and identified 825 genes annotated by NCBI as pseudogenes. We excluded these genes from all analyses, together with any gene whose ortholog was flagged as pseudogene in the *Mus musculus* annotation (GENCODE v15).

### t-SNE / PCA / distance-based clustering

To investigate the main axes of variation in gene expression in our dataset, we used three different approaches. Prior to these analyses, we log-transformed and normalized count data using the regularized log-transformation function (rlog) implemented in *DESeq2* (Love, Huber, and Anders 2014). First, we used t-distributed stochastic neighbor embedding (t-SNE) implemented in *Rtsne* (Krijthe 2015). Second, we performed a principal component analysis (PCA) on centered and scaled data. Finally, we performed hierarchical clustering based on Euclidean distance. A single F_1_ hybrid hypothalamus sample was identified as an outlier in all three approaches, and therefore we excluded this sample from all downstream analyses.

### Differential expression (DE) analyses and multivariate adaptive shrinkage

We tested for differential gene expression (1) between sexes within species and (2) between the two species independent of sex. Briefly, for each comparison, we used *DESeq2* to obtain log_2_ fold changes (effect size estimates) together with their associated per-gene standard errors and then used multivariate adaptive shrinkage (MASH) (Urbut et al. 2019) to adjust these effect size estimates and assess their statistical significance (see Suppl. Materials A for details).

To test if the number of DE genes was biased towards a species or sex, we performed exact binomial tests in *R*, assuming an equal probability of success (p=0.5) and correcting for multiple testing using the Bonferroni-Holm approach. In analyses comparing the concordance between individual subregion and whole brain differential expression calls, we used *DEseq2* and called statistically differentially expressed genes (p < 0.05) in each subregion and the whole brain separately, correcting for multiple testing using the Benjamini-Hochberg procedure.

### Gene regulation analyses

For these analyses, we estimated gene expression with *bam2hits* and *MMSEQ* (Turro et al. 2011). We assigned genes to five gene regulatory classes based on the individual expression estimates in both parental species and alleles in F_1_ hybrids: “conserved”, “compensatory”, “*cis*”, “*trans*”, and “*cis* & *trans*” using *MMDIFF* (Turro, Astle, and Tavaré 2014). To compare the relative support (i.e., posterior likelihood) for each of the five models for any given gene, we used a polytomous model comparison approach (see Turro et al. 2014 for details) using a flat prior probability of 0.2 for each model. Design matrices for all models included sex as a covariate to account for potential confounding effects. We performed all gene regulation analyses separately for each subregion. We evaluated the performance and robustness of our model selection approach using a shuffling and a bootstrapping approach (see Suppl. Materials B). Based on these analyses, we decided to use a posterior probability threshold of 0.75, which resulted in almost no false positives in the shuffled controls and more than 95% concordant assignments in our bootstrap approach (Fig. S3).

After statistical assignment via model comparisons, we further subdivided genes in the “*cis* & *trans*” class into the “*cis* + *trans*” and “*cis* x *trans*” classes (Landry et al. 2005) based on the ratio of weighted log-fold changes (Shen, Turro, and Corbo 2014) in the parental species to weighted log-fold changes of alleles in F_1_ hybrids.

Because all male F_1_ hybrids carry exclusively the *P. maniculatus* X chromosome, we did not consider genes on the X chromosome in the gene regulation analyses. Furthermore, we only kept genes expressed at a minimum μ (equivalent to RPKM; Turro et al. 2011) of one in at least three animals. We omitted between 2–21 genes per subregion due to model convergence issues, together with an additional 138 genes that are subject to genomic imprinting (Perez et al. 2015). Finally, we performed clustering of categorical data in heatmaps of inter-subregion regulation change based on Gower distance and a divisive algorithm.

### Gene enrichment analyses

We tested if the function of sets of genes of interest were overrepresented with *WebGestaltR* (Liao et al. 2019). Applying a false discovery rate of 0.05 and using *Mus musculus* as a reference, we queried the non-redundant gene ontology databases as well as the Mammalian Phenotype Ontology, the KEGG, Panther, and Reactome pathway databases. If indicated, we reduced the redundancy of results to a maximum of 10 sets by applying *WebGestaltR’s* weighted set cover function. As background, we only retained genes that were expressed in our data set and passed the same filtering thresholds as the genes of interest (foreground) on an analysis-by-analysis basis.

## Results

### Main axes of variation in gene expression separate subregions and species

As a first step, we characterized overall patterns of gene expression across subregions in both species. We also characterized gene expression in whole brain samples for comparison with subregions. Overall, we found 16,078 genes (excluding pseudogenes) are expressed in the brain, using a threshold of at least one count per million reads (CPM) in at least three samples in any given subregion. t-distributed stochastic neighbor embedding (t-SNE) showed that samples cluster consistently both by subregion and species (Fig. 1C). This finding was also supported by a principal component analysis (PCA) (Fig. S2A): three axes of the PCA explain most of the variation between whole brain and subregions (PC1, 17.85%), across subregions (PC2, 14.95%), and between cerebellum and all other subregions (PC4, 12.08%), and an additional axis (PC3) explains variation between species (13.59%). Finally, the whole brain and cerebellum gene expression profiles were the most distinct in a hierarchical, distance-based clustering approach (Fig. S2B). Together, these results show that each brain subregion exhibits a distinct transcriptomic profile, but also that there are interspecific gene expression differences that persist across subregions.

### Most genes are expressed in all subregions while few are subregion-specific

Given that variation in gene expression was primarily driven by differences among subregions, we next evaluated the proportion of genes with subregion-specific expression patterns. We found that each individual subregion expressed between 13,755 and 14,429 genes (of the 16,078 genes that met our expression threshold). Of these, 12,769 genes (82.03% of genes included in this analysis) were expressed in all ten subregions and only 623 genes (4.00%) were expressed exclusively in one subregion (Fig. 1D). The top three most enriched terms of these privately expressed genes represented specialized functions: “Hormone ligand-binding receptors”, “Glycoprotein hormones”, and “Peptide hormone biosynthesis” (all three in the Reactome pathway database). The highest number of privately expressed genes was found in the cerebellum and the lowest in the amygdala (Fig. 1E). An additional 511 genes were captured only in the whole brain samples and are likely regionally restricted to areas not captured by our subregion sampling. The small number of region-specific genes, together with the regionalized transcriptional profiles (Fig. 1C), suggests that specialized functions arise from the coordination of many genes acting in concert rather than through single genes.

### Whole brain samples only partially capture subregion-specific expression divergence between species

Because comparative studies often use whole brain dissections rather than specific subregions, we investigated the extent to which region-specific differential expression between species is reflected in a sample generated from the entire brain. In total, we found 12,624 genes (78.5% of the 16,078 genes included in this analysis) were significantly differentially expressed (DE) (local false sign rate < 0.05) in at least one subregion or the whole brain, and 3,462 genes (21.5%) were significantly DE in all subregions and the whole brain. We found that 58.4–63.3% of DE calls were significant (FDR < 0.05) and in the same direction between the whole brain and any single subregion (Fig. S4). In 10.0–19.5% of cases, there was a significant difference in the respective subregion but not the whole brain, which appears to be driven by localized DE that is not detectable (too diluted) in a whole brain sample (see Supp. Materials C for additional analyses). We also found 16.2–22.7% of cases where a DE call was in the whole brain but not in any given subregion, which likely reflects localized DE in a region not sampled in our subregion dissections. Only in 0.8–2.2% cases were mismatches due to a “sign flip” where the direction of expression differences (log_2_ fold changes) was inverted yet significant in both the whole brain and a specific subregion. Taken together, these results suggest that whole brain samples capture most, but not all, of the subregion-specific DE genes.

### Differential gene expression is pervasive between species

Given that each subregion shows a distinct transcriptomic profile, we next compared differential gene expression (DE) between the two species across subregions. Instead of analyzing each subregion separately, we used multivariate adaptive shrinkage to account for correlations in effect sizes among subregions and to improve power (Urbut et al. 2019). Within subregions, the number of DE genes ranged from 52.0–59.3% of expressed genes, with no bias towards either species (exact binomial test; all p_adj_ > 0.05) (Fig. 2A). To focus on genes that most likely have biologically relevant levels of differential expression, we applied a conservative effect-size filter of a minimum two-fold change in DE between species. This filter reduced the number to 2,919 genes (18.2% of all expressed genes). Most of these genes – especially the most differentially expressed ones – were consistently differentially expressed in the same direction across brain regions (Fig. 2B). However, 628 of these (21.5% of the 2,919 genes) showed a sign flip in log_2_ fold changes across subregions (Fig. 2B), indicating considerable subregion-specific variation in differential expression. To quantitatively assess the correlations in species-biased gene expression among subregions and the whole brain, we calculated the proportion of sharing by magnitude (i.e., the proportion of genes significantly DE in at least one of the two tested subregions that exhibited a log_2_ fold change within a factor of 0.5 of each other) among all pairwise comparisons. The proportion of sharing ranged from 0.51 to 0.86 (Fig. 2C). The highest correlations were found among subregions such as midbrain and hypothalamus (0.86) and amygdala and septum (0.85). By contrast, correlations between subregions and the whole brain showed the lowest levels of pairwise sharing with any of the subregions, followed by the cerebellum. These patterns are largely consistent with the developmental origin of each subregion. Together, these data show widespread DE between all subregions of the brains of these closely related species, but that the identity of DE genes varies across subregions.

**Fig. 2.**
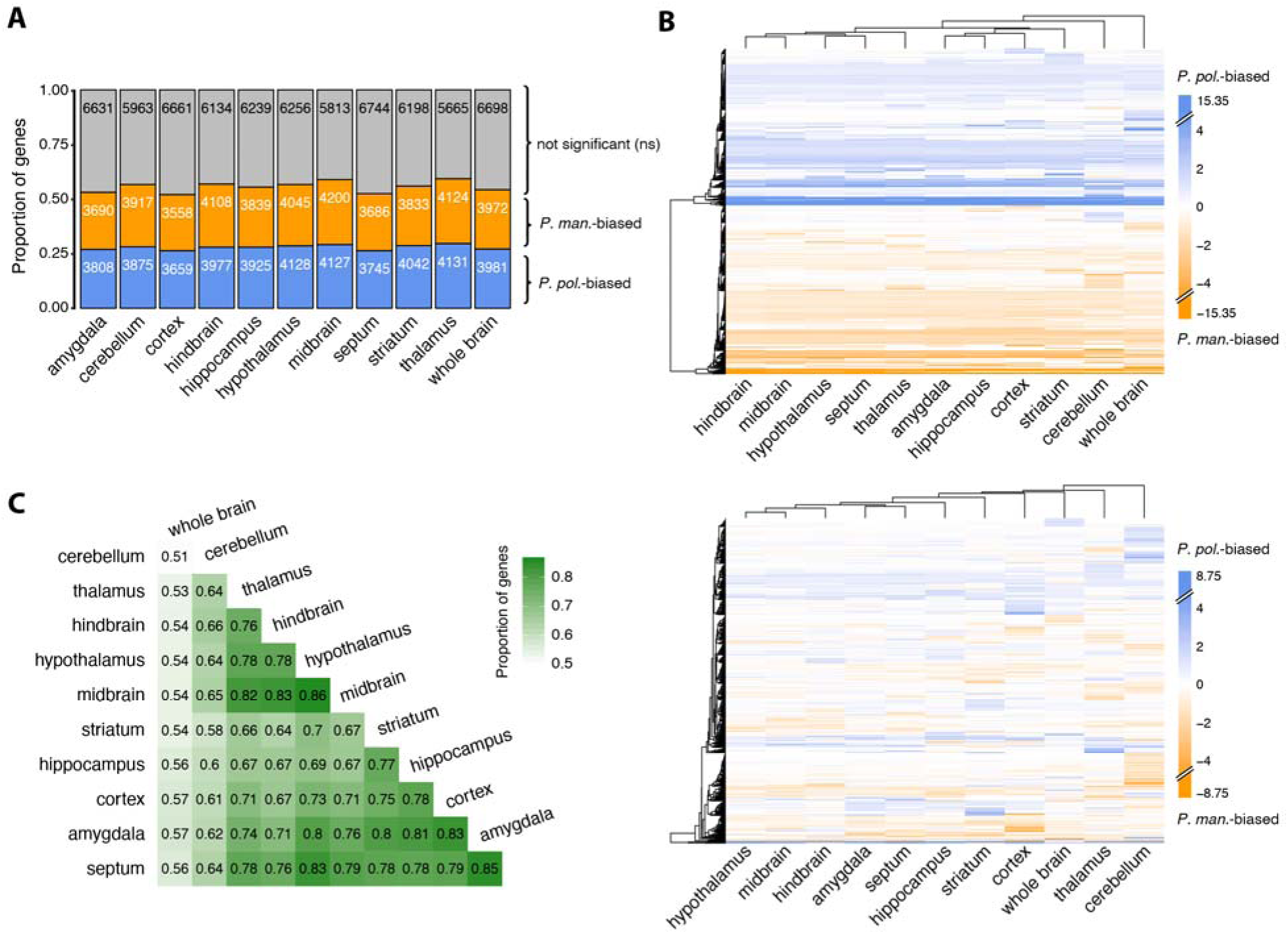
Species-biased gene expression across subregions and the whole brain. (**A**) Proportion and number of significantly differentially expressed (DE) genes biased towards either species across 10 subregions and the whole brain. (**B**) Adjusted log_2_ fold change (LFC) estimates (after multivariate adaptive shrinkage) across subregions and the whole brain of 2,919 genes that are significantly differentially expressed and exhibited a log_2_ fold change > 1 in at least one subregion (top); subset of 628 genes that show a sign flip in LFC estimates across subregions (bottom). Note that maximum color intensity was cut off at ± 5 LFC. Maximum and minimum LFC estimates are indicated in the legend. (**C**) Proportion of significantly DE genes in any pairwise comparison that exhibited a shared signal, defined as exhibiting a log_2_ fold change in the same direction within a factor of 0.5 of each other.

### Lack of support for universal transcriptomic mechanism underlying monogamy

Previous work has compared whole brain gene expression between pairs of promiscuous and monogamous species across four major vertebrate clades and proposed a set of 41 candidate genes associated with monogamous mating systems (Young et al. 2019). Given the difference in mating systems in our two focal species (*P. maniculatus* = promiscuous; *P. polionotus* = monogamous), we wanted to test whether the same set of genes are identified as DE in our data. Of the candidate genes identified in Young et al. 2019, 38 were present in our *Peromyscus* transcriptome and 30 of the 38 were also DE across our focal species. Of these 30 genes, one was reported to be up-regulated in promiscuous species, while the remaining 29 were up-regulated in monogamous species. In our data, however, only 15 of 30 (50%) were DE in the same direction (in any subregion) (Fig. S5), consistent with the number of direction-matching DE genes expected by chance. Thus, we do not find evidence for the proposed universal gene set underlying monogamy in our data.

### Sex-biased gene expression is often species-specific

In addition to mating system differences, the two focal *Peromyscus* species also differ in their degree of sexually dimorphic reproductive behaviors, including parental care, which is performed by females of both species but, in males, is largely restricted to *P. polionotus* and limited in *P. maniculatus* (Bendesky et al. 2017). Thus, we were also interested in identifying genes with a species-specific sex bias. We note that our whole brain samples were collected exclusively from males and therefore not suited for these analyses.

In total, we found 461 significantly DE genes between females and males in at least one of the 20 species and subregion combinations, corresponding to 3.0% of all 15,503 genes in this analysis. While 203 out the 461 genes were shared (i.e., showed a sex-bias in at least one subregion) between the two species, 150 genes were sex-biased only in *P. polionotus* and 108 only in *P. maniculatus*. The number of sex-biased genes ranged from 118–205 with a significantly higher number of male-biased than female-biased genes in seven out of 20 species-subregion comparisons (exact binomial test; p_adj_ < 0.05; Bonferroni-Holm correction). Only the hippocampus of *P. polionotus* exhibited the reverse pattern of a significant excess of female-biased genes over male-biased genes (Fig. 3A). In addition, the total number of sex-biased genes (male-biased plus female-biased) was higher in *P. polionotus* than *P. maniculatu*s in the cerebellum and hippocampus, and vice versa in the septum (exact binomial test; p_adj_ < 0.05; Bonferroni-Holm correction) (Fig. 3A). Per subregion, between 28.4–41.3% of sex-biased genes were shared between the two species (Fig. 3B). The log_2_ fold change distributions of these shared sex-biased genes were overall very similar in both species across subregions (Fig. 3C). Thus, we detected no consistent quantitative differences in sex-biased genes, neither in overall number nor in expression magnitude (log fold changes in shared genes) between the two species.

**Fig. 3.**
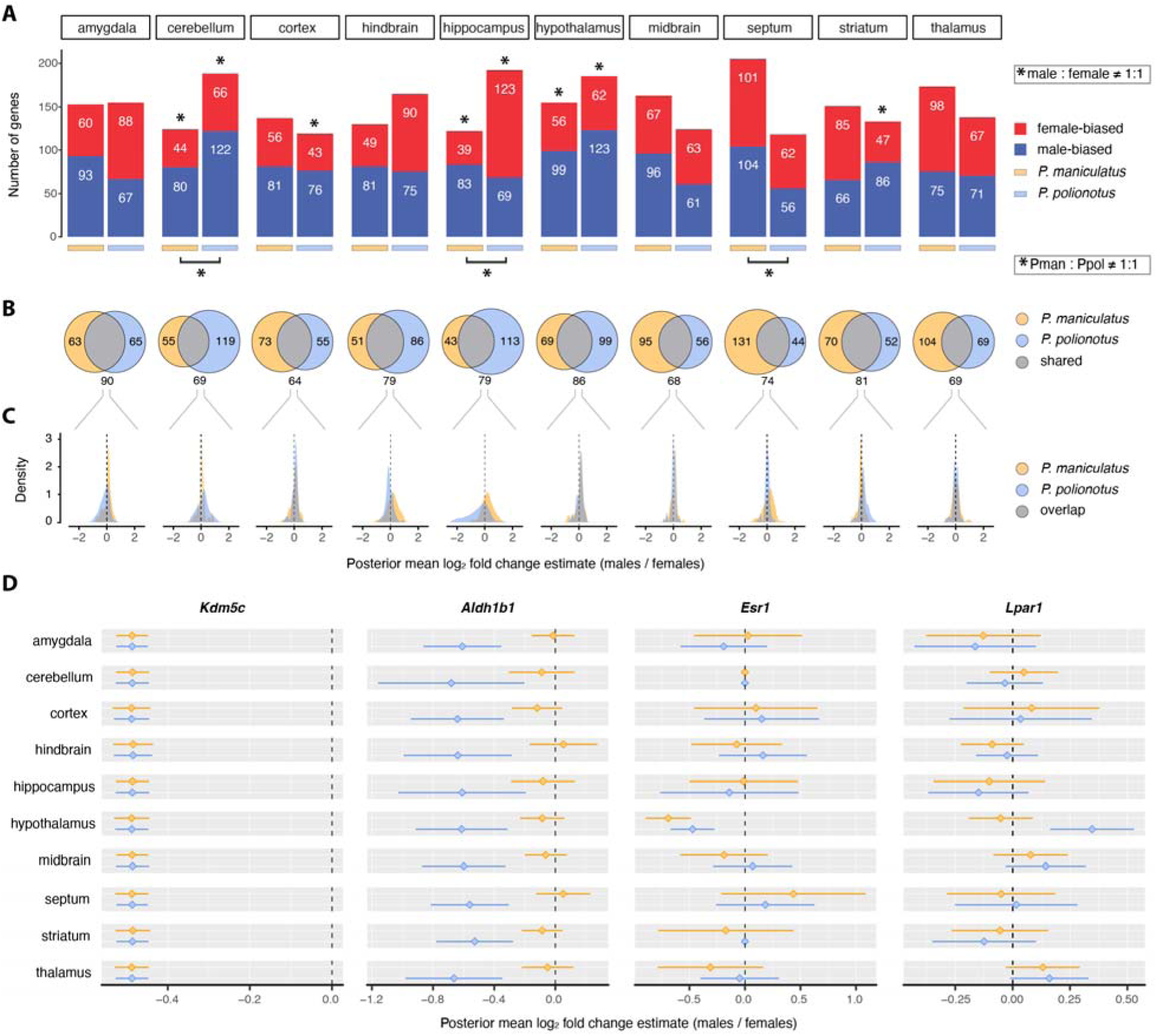
Shared and species-specific sex-biased gene expression. (**A**) Number of significantly differentially expressed female-or male-biased genes by species and subregion. Asterisks indicate statistically significant differences between males and females within species (on top of bars) and between species (below bars) based on binomial tests after Bonferroni-Holm family-wise error rate correction (p<0.05). (**B**) Number of shared and species-specific sex-biased genes. (**C**) Distribution of log_2_ fold changes for shared sex-biased genes in both species. (**D**) Exemplary sex-biased genes, either sex-biased in both species across all subregions (*Kdm5c*), only one species across all subregions (*Aldh1b1*), in both species in only one specific subregion (*Esr1*), or one species in only one specific subregion (*Lpar1*). Diamonds and error bars indicate posterior mean log_2_ fold change estimates (after multivariate adaptive shrinkage) with 95% confidence intervals (CI) (mean ± 1.96 SD). Estimates of zero with a CI of zero indicate genes that were not expressed in a specific subregion.

Many previously known sex-biased genes were differentially expressed between females and males in both species across all subregions (e.g., the X-linked Lysine-specific demethylase 5C [Kdm5c]) (Naqvi et al. 2019), or only in specific subregions (e.g., estrogen receptor 1 [Esr1] in the hypothalamus) (Xu et al. 2012) (Fig. 3D). However, we also detected several genes that were sex-biased in only one species and not the other in both the monogamous and the promiscuous species. Some of the genes were consistently sex-biased in all subregions of one species (e.g., aldehyde dehydrogenase 1 family, member B1 [Aldh1b1] in *P. polionotus*), whereas others were so in only specific subregions (e.g., Lysophosphatidic acid receptor [Lpar1] in the hypothalamus of *P. polionotus*) (Fig. 3D). These data emphasize that sex-specific DE is only partially shared between species.

### Differential gene expression between species is mostly due to *cis*-regulatory divergence

By leveraging allele-specific expression patterns in F_1_ hybrids (Fig. S6A), we investigated the regulatory bases underlying gene expression divergence between species. *Cis*-regulatory changes are often presumed to have local, tissue-specific effects and could therefore be of particular importance in the evolution of gene expression in a heterogenous organ like the brain compared to *trans*-regulatory changes, which potentially have pleiotropic, broad effects (Signor and Nuzhdin 2018). In addition, quantifying the impact of regulatory changes (*cis* vs. *trans*) can inform the relative roles of selection (stabilizing and divergent) versus neutral evolution driving gene expression differences between species (see below).

We assigned genes to the five following regulatory classes using a Bayesian regression framework followed by polytomous model comparisons: “conserved”, “compensatory”, purely “*cis*”, purely “*trans*”, and “*cis* & *trans*” (Fig. S6B). The “*cis* & *trans*” class was further subdivided *post hoc*: *cis*-and *trans*-regulatory effects that act synergistically in driving gene expression divergence were classified as “*cis* + *trans*” and *cis*- and *trans*-regulatory effects that act in opposite directions were classified as “*cis* x *trans*” (Fig. 4A). No expression difference between either the parental species or alleles in F_1_ hybrid is consistent with conserved gene expression regulation, whereas no expression difference between parental species but differential expression of alleles in F_1_ hybrid is consistent with compensatory gene regulation. Both classes together, “conserved” and “compensatory”, comprise non-differentially expressed genes between species and comprise 41.8–54.2% of all genes across subregions (Fig. 4C). While the lack of regulatory divergence in the conserved class could be due to stabilizing selection or neutral evolution, compensatory gene regulation is more consistent with stabilizing selection (Landry et al. 2005).

**Fig. 4.**
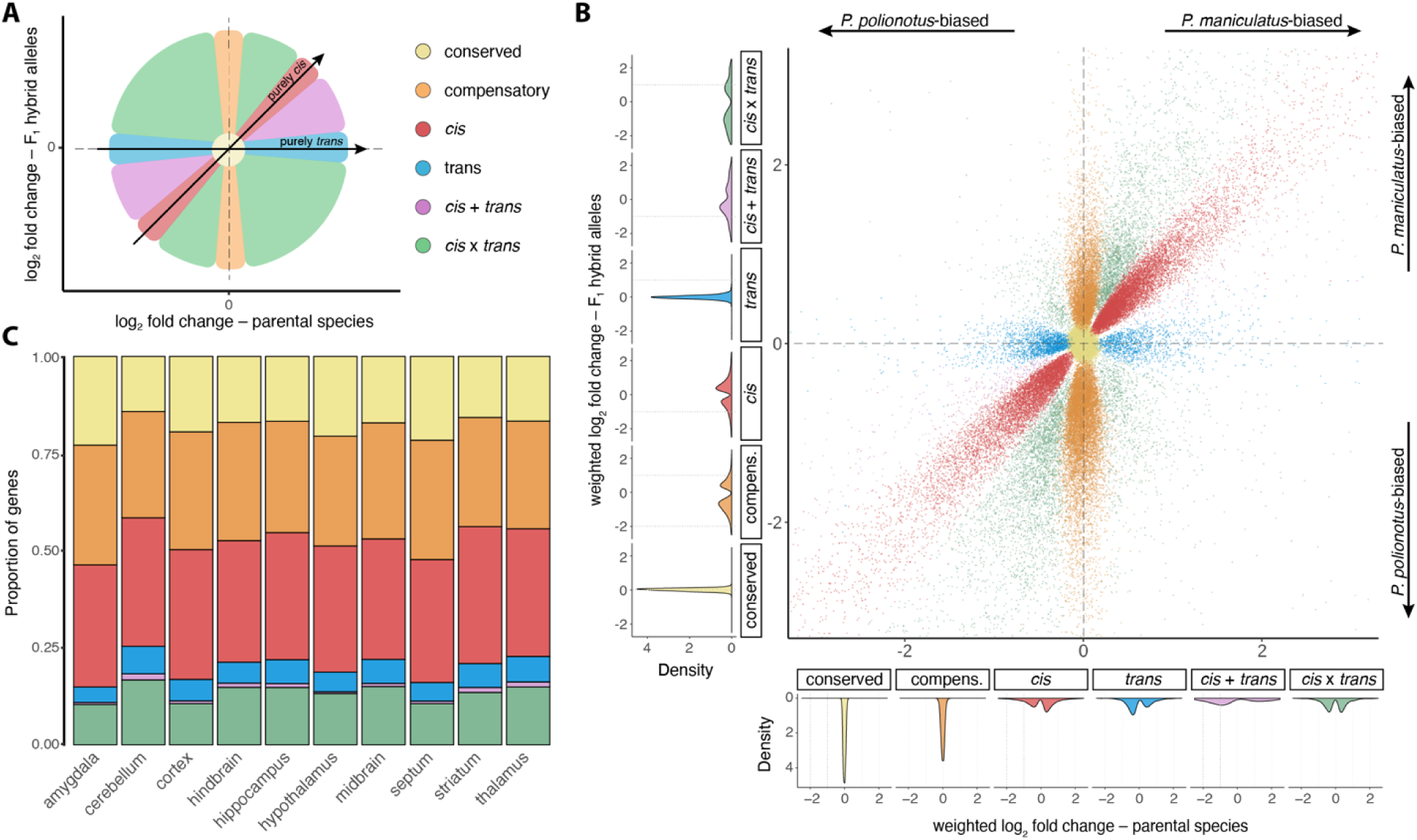
Assignment of genes to gene regulatory classes. (**A**) Schematic of gene regulatory classes in relation to expression ratios in parental species versus F_1_ hybrid alleles. (**B**) Empirically determined weighted gene expression log_2_ fold changes in parental species versus log_2_ fold changes of alleles in F_1_ hybrids across the entire data set. Each dot is a single gene color-coded by its inferred gene regulatory class (applying posterior probability cutoff of 0.75). Side-panels provide distribution of weighted log_2_ fold change estimates for parental species (bottom) and alleles in F_1_ hybrids (left) by gene regulatory class. Genes with stark expression differences (log_2_ fold change of more than plus/minus three) were omitted to enhance readability (see Fig. S7A for full distribution). (**C**) Proportion of genes assigned to different gene regulatory classes across subregions. Genes that could not be assigned (at this threshold) were omitted for clarity from all panels (see Fig. S4E for proportion of unassigned genes).

The other regulatory classes, “*cis*”, “*trans*”, “*cis* + *trans*” and “*cis* x *trans*”, which encompass genes that show expression differences in the parental species, account together for 45.8–58.2% of genes across subregions (Fig. 4C) – a range close to the 52.0–59.3% observed in our DE analysis using a different methodological approach and considering only the parental species (Fig. 2A). While most of these genes exhibited modest expression differences (log_2_ fold change [LFC] > -3 and < 3) (Fig. 4B), most of the large expression differences (LFC < -3 or > 3) were due purely to *cis*-regulatory divergence (Fig. S7A). This pattern was consistent across individual subregions (Fig. S7B-D). Purely *cis*-regulatory divergence was also the most common regulatory class underlying expression differences with 30.7–35.0% compared to only 4.0–7.2% purely *trans*-regulatory divergence across subregions (Fig. 4C). While DE due to purely *cis*- or purely *trans*-regulatory divergence could be due to divergent selection or neutral evolution, an accumulation of mutations driving gene expression divergence between species in the same direction (“*cis* + *trans*”) and is more consistent with divergent selection (Orr 1998; Fraser, Moses, and Schadt 2010; Verta and Jones 2019), and accounted for 0.4–1.6% of genes. Taken together, these analyses demonstrate that most of gene expression divergence between these two species is due to *cis*-regulatory differences.

To quantitatively assess whether expression in F_1_ hybrids is regulated in an additive, dominant, or transgressive (over- or under-dominant) way (Gibson et al. 2004; Landry et al. 2005), we also classified genes in an orthologous manner by their overall expression levels in parental species versus F_1_ hybrids (Fig. S8A). Consistent with a study in three-spined stickleback fish (Verta and Jones 2019), we found genes in the “*cis*” and “*cis* + *trans*” classes to be expressed in a slightly more additive way than genes in the “*trans*” and “*cis* x *trans*” classes (Fig. S8B). The “*cis* x *trans*” class also contained many genes that showed transgressive expression patterns (Fig. S8B); such misexpression of genes in hybrids due to unbalanced *cis*- and *trans*-regulatory mechanism could contribute to hybrid incompatibilities and thereby speciation or species maintenance (McGirr and Martin 2019).

### High degree of modularity in gene regulation across the brain

Having quantified the relative sizes of gene regulatory classes among subregions, we were motivated to assess how consistent gene regulation was for any given gene across subregions. In other words, how frequently is a gene assigned to the same gene regulatory class across all ten subregions. In total, 6,600 genes (43.1% of the 15,308 genes included in this analysis after filtering; see Methods for details) were exclusively assigned to a single regulatory class (Fig. 5A). Out of these, 1,696 genes could only be assigned unambiguously in a single subregion. Disregarding these genes leaves 4,904 genes (32.0%) that could be assigned in more than one subregion and were consistently assigned to the same regulatory class. Interestingly, just as many genes, 4,975 (32.5%), were assigned to two different regulatory classes. A smaller, but considerable, number of 1,171 (7.6%) genes was assigned to three different classes, and a fraction of genes, 109 (0.7%) and 6 (<0.1%), were even assigned to four or five different classes, respectively (Fig. 5A). The remaining 2,447 genes (16.0%) could not be assigned to any regulatory class in any single subregion at the applied posterior probability threshold of 0.75.

**Fig. 5.**
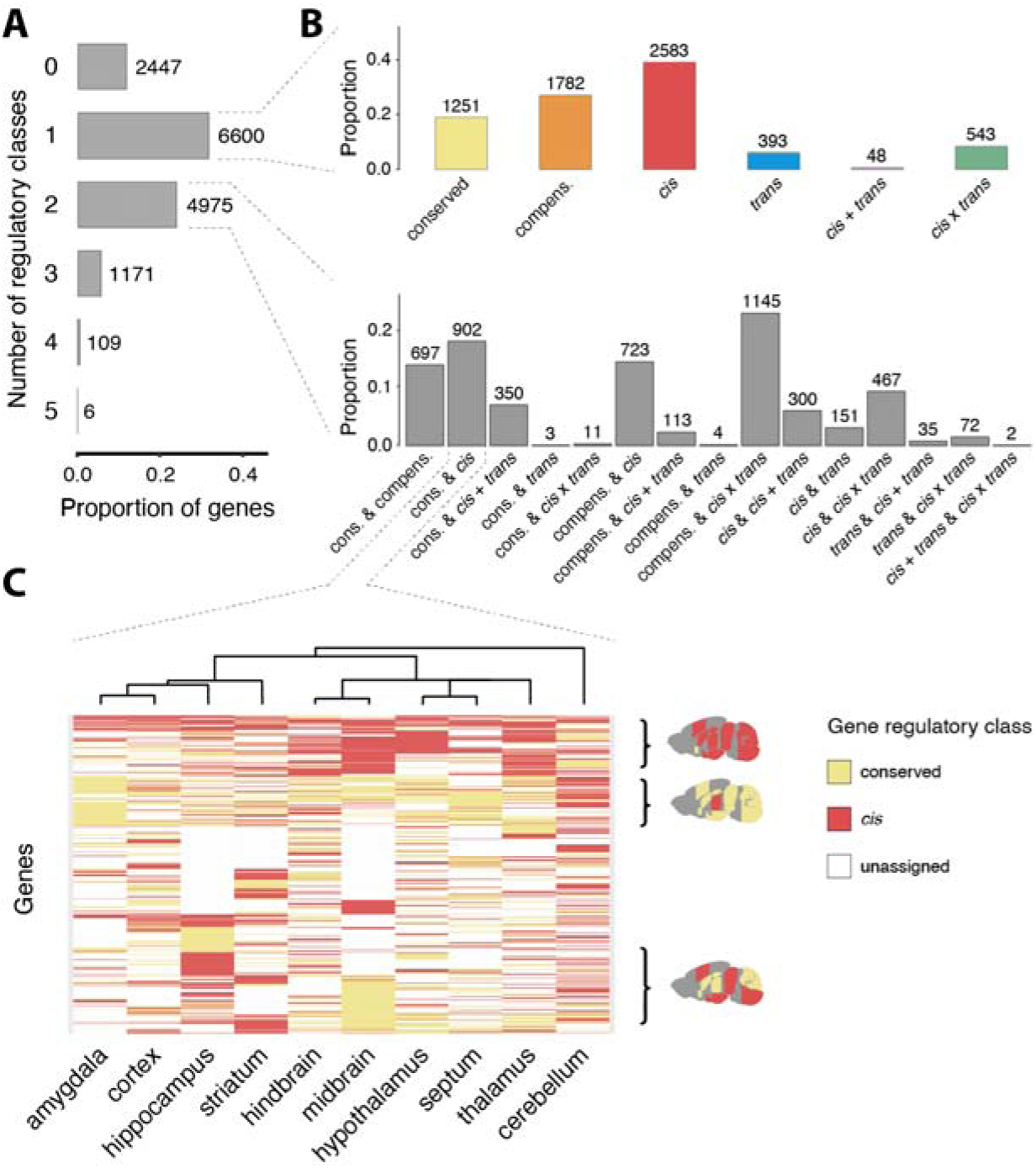
Shared versus modular gene regulation across subregions of the brain. (**A**) Proportion and number of genes that were assigned to either none, only one, or up to 5 different regulatory classes across subregions. (**B**) Breakdown of the proportion of genes consistently assigned to one of the 6 classes (top) or one of the 15 two-class combinations for genes consistently assigned to only one or exactly two different classes (bottom). (**C**) Regulatory class assignment across subregions of all genes (rows) that exhibited both conserved gene expression and differential expression due to *cis*-regulatory divergence in at least one subregion. Some genes showed differential expression due to *cis*-regulatory divergence in all but one subregion (where gene expression was conserved) and vice versa or a mix of regulation patterns, indicated by brain icons on the right. Subregions and genes were clustered based on Gower distance. Genes that could not be assigned at the applied posterior probability threshold of 0.75 in any given subregion are shown in white.

Considering only genes that were consistently assigned to one class, the “*cis*” class was again the largest, followed by the “compensatory”, “conserved”, “*cis* x *trans*”, “*trans*”, and “*cis* + *trans*” classes (Fig. 5B), akin to the frequencies observed when we considered subregions separately (Fig. 4C). Genes consistently assigned to the “compensatory” and “conserved” classes might be under strong selective constraints and, indeed, were enriched for the terms “preweaning lethality” and “abnormal survival” (Mammalian Phenotype Ontology database). Genes consistently in the “*cis* x *trans*” class, in which *cis*- and *trans*-regulatory differences act in a compensatory but not completely balanced way, are likely to also be under selective constraints and were enriched for terms involved in nuclear processes, including “single-stranded RNA binding” and “translation factor activity, RNA binding” (Molecular Function GO database), and “metabolism of RNA” (Reactome pathway). In contrast, genes assigned to the “*cis*” class were enriched for the term “receptor regulator activity” (Molecular Function GO database). The two smallest classes, “*trans*” and “*cis* + *trans*” were not statistically significantly enriched for any terms.

Of the genes that were assigned to two different classes, the combination of “compensatory” and “*cis* x *trans*” was the largest (Fig. 5B), which is not unexpected given that the “compensatory” class can be considered a special (completely balanced) case of “*cis* x *trans*” regulatory divergence. The second largest combination was “conserved” together with “*cis*”. This combination is interesting in that it contains genes that are differentially expressed between the parental species in some subregions but not others, which implies that regulation is mediated by *cis*-regulatory divergence that is acted upon by subregion-specific transcription factors. Focusing on these “conserved & *cis*” genes, we found a broad spectrum of regulation patterns, ranging from genes being affected by *cis*-regulatory divergence in most subregions to the opposite extreme of genes whose regulation is conserved in almost all subregions (Fig. 5C). Based on the (categorical) assignment of genes in this subset the cerebellum was again the most distinct subregion, standing apart from one cluster comprising the amygdala, cortex, hippocampus, and striatum and another cluster comprising the hindbrain, midbrain, hypothalamus, septum, and thalamus (Fig. 5C). In conclusion, we find that the evolution of gene expression divergence between the two focal species happens to a considerable extent through subregion-specific mechanisms across the brain.

## Discussion

The evolution of gene expression in the brain and how it contributes to the evolution of behavior remains poorly understood. Here, we focused on two closely related yet behaviorally divergent species and compared gene expression across brain regions, between species, and between sexes as a first step towards understanding how their brain transcriptomes evolve. We find that slightly more of the variation in gene expression is across brain subregions rather than between species, echoing similar patterns found in comparative gene expression studies of different tissues (Brawand et al. 2011; Merkin et al. 2012). This suggests an evolutionarily conserved transcriptional program for each subregion of the brain that may reflect the specialized functions of each subregion. Nonetheless, subregion transcriptomes form clusters that reflect their developmental history. Cortex, hippocampus, amygdala, and striatum form one cluster, while midbrain, hypothalamus, and thalamus form another, with septum falling in between. Cerebellum is the most distinct subregion of all. The clustering of hindbrain with midbrain in our data may reflect difficulties in dissecting pure subregion samples or batch effects. These associations are also reflected in differential expression (DE) patterns between species, implying the presence of shared regulatory programs across subregions.

Importantly, even though our data imply conserved transcriptional profiles for each subregion, we also detect pervasive DE between species: most of the brain transcriptome is differentially expressed in at least one subregion. This is in line with expectations, given extensive variation in gene expression observed even among strains of a single species (Nadler et al. 2006). Given that our two focal species differ in their mating system, we were interested in testing whether our data would support previous work that proposed a set of candidate genes robustly associated with monogamous mating systems across vertebrates (Young et al. 2019). Our results do not provide evidence for this universal transcriptomic signature, consistent with a re-analysis of Young et al. 2019 that did not find evidence of transcriptome-wide parallel evolution in the repeated evolution of monogamy (Jiang and Zhang 2019). We note, however, that behavioral transitions may induce gene expression differences in the brain (Hu et al. 2022; Ray et al. 2016). Our analyses were performed in virgin animals and may therefore not have captured the DE changes initiated by reproductive events.

Sex-specific behaviors such as parental care behavior are also known to correlate with mating system. In *Peromyscus* deer mice, large differences in parental behavior are observed between our two focal species, but the differences are more pronounced in fathers than mothers (Bendesky et al. 2017). Motivated by this observation, we looked at differential expression across the sexes, but did not find obvious patterns of increased sex bias in one species or the other. However, we note that our bulk RNA-sequencing strategy may not provide the spatial and cellular resolution to uncover functionally important sexually dimorphic nuclei (Xu et al. 2012; Kim et al. 2019). We believe this avenue of research may be of interest for future studies.

Finally, we sought to understand the regulatory mechanisms underlying the evolution of gene expression, leveraging F_1_ hybrids to differentiate *cis*- and *trans*-regulatory factors and assign genes to gene regulatory classes (e.g., *cis*, *trans*, *cis* & *trans,* etc.). Almost half the transcriptome is conserved in expression due to lack of regulatory differences or compensatory gene regulation. This is evidence that a large portion of the transcriptome is under stabilizing selection and explains the conserved transcriptional programs we found for each brain subregion. Of the genes differentially expressed between species, we found that the majority is driven by *cis*-regulatory factors. The predominance of *cis*-regulatory changes in expression evolution between species has been found repeatedly across many animal taxa including Mexican cavefish (Leclercq et al. 2022), stickleback (Verta and Jones 2019), *Mus* (Reuveni et al. 2018), and *Drosophila* (Benowitz et al. 2020). These observations and our data are consistent with the prediction that selection on gene expression would be driven primarily from *cis*-rather than *trans*-regulatory evolution due to the reduced pleiotropy of *cis*-acting changes (which tend to occur in promoters or enhancers of target genes) compared to *trans*-regulatory changes (which tend to affect transcription factors that have downstream effects on large networks of genes) (Signor and Nuzhdin 2018). While many genes were assigned exclusively to a single regulatory class, we find over 6,000 genes that are evolving via two or more different regulatory classes across subregions of the brain, suggesting that region-specific regulatory evolution is widespread. This set of genes may include those that are undergoing region-specific selection and may be of interest for future follow-up studies interested in behaviors likely mediate through region-specific gene functions.

Our study highlights the complexity of gene expression evolution in the brain. Even between closely related species, we observe widespread DE. This DE is driven by a combination of correlated changes across several or all subregions and region-specific regulatory changes, predominantly in *cis*. We hypothesize that this generates modularity, which may be key in providing the evolutionary substrate for the myriad of behavioral phenotypes displayed by animals. Having extensively characterized gene expression differences and their regulatory basis across subregions, our study lays the groundwork for future investigations that seek to understand the mechanistic basis of behavioral evolution, in deer mice and beyond.

## Acknowledgments

All computational analyses were performed on the Harvard University’s Faculty of Arts & Sciences (FAS) research computing cluster.

## Author Contributions

CLL, CH, AB, and HEH conceived of the study. FB, KT, and AB set up crosses to generate F_1_ hybrids. CLL, CH, NLB, and FB performed tissue dissections. KT generated libraries for sequencing. AFK, JC, CLL, and JML analyzed data. CLL, CH, AB, and HEH acquired funding. AFK, JC, and HEH wrote the manuscript with contributions from CLL, CH, and KT. All authors approved of the final version of the manuscript.

## Data Accessibility Statement

Raw sequence reads and gene expression estimates will be deposited on the SRA and GEO. Custom code to reproduce these results will be uploaded to GitHub.

