## Supplementary Text for "Evolution of gene expression across brain regions in behaviorally divergent deer mice"

### **Supplementary materials**

(A) Multivariate adaptive shrinkage

Multivariate adaptive shrinkage offers increased power over a subregion-by-subregion approach and reduces the chances of underestimating shared expression differences due to simple threshold-based effects, that is, a gene with a similar effect size in two subregions barely passing the significance threshold in one but not the other subregion. We tested for differential gene expression between sexes in all 10 subregions, but not the whole brain samples, since the latter were collected from males only. Consequently, we investigated differential gene expression between sexes in 20 different species-subregion combinations (10 subregions in both species) and between species in 11 different subregions plus whole brain comparisons. We omitted F_1_ hybrid animals from these analyses. For each condition – keeping only genes expressed at a minimum of one count per million in at least three animals – we fitted a model in *DESeq2* including only sex for the sex DE analyses and species, sex, as well as a sex-species interaction in the species DE analyses to account for confounding effects. We normalized count data using the implemented median of ratios approach (Anders & Huber, 2010). We then used the estimated fold changes and standard errors as input for *mashr* (Urbut et al. 2019). Following (Naqvi et al., 2019), we set missing values (i.e., genes not expressed in certain conditions) to zero with a standard error of 1,000. *Mashr* can learn patterns from the data and use them to improve effect size estimates (shrinkage) using an empirical Bayes method. Running *mashr* is a multi-step approach. First, we used the implemented *estimate_null_correlation_simple* function to estimate correlations among conditions. Next, we calculated canonical covariance matrices and data-driven (i.e., learned) covariance matrices. While obtaining the former is straightforward using the *cov_canonical* function, obtaining the latter is a multi-step procedure. Following (Naqvi et al., 2019) (see also https://stephenslab.github.io/mashr/articles/intro_mash_dd.html), our approach consisted of the following four steps: (i) extract genes with strong signals from each condition separately (local false sign rate < 0.05; Stephens 2017) in a condition-by-condition manner using the *mash_1by1* function; (ii) perform a PCA on these genes using the *cov_pca* function generating six covariance matrices (one for each of the first five PCs and one based on all of them); (iii) perform a sparse factor analysis (SFA; Engelhardt and Stephens 2010) on a matrix of Z-scores (fold change estimates divided by standard errors) assuming five clusters and convert factors to covariance matrices with *cov_from_factors*; and (iv) use the PCA- and SFA-derived covariance matrices to initialize the implemented extreme deconvolution algorithm with *cov_ed* and obtain the final data-driven covariance matrices. Finally, we obtained the shrinked fold change estimates running the main *mash* function using both the canonical and data-driven covariance matrices (the log-likelihood of the model using both together was higher than using each separately). We ran the whole procedure separately for sex-bias (males vs. females in 20 conditions / subregions in both species) and species-bias (*P. maniculatus* vs. *P. polionotus* in 11 conditions / subregions and whole brain) analyses. We called differentially expressed genes in each condition at a local false sign rate < 0.05, extracted the pairwise proportion of sharing using *mashr’s* *get_pairwise_sharing* function, and visualized the results with the *corrplot()* R package. To avoid biasing the inferred extent of sharing, we set effect sizes (log_2_ fold change) of genes that had missing values in certain subregions (due to sub-threshold or no expression) prior to multivariate adaptive shrinkage to zero immediately afterwards (and prior to downstream analyses).

### (B) Shuffling and Bootstrapping

First, we assigned half of the samples and F_1_ hybrid alleles of one species to the other species and vice versa. The expectation of this shuffling design is that any expression differences should average out and the most supported model should therefore be the “conserved” one. Second, to test the influence of individual (outlier) samples, we used a non-parametric bootstrap approach. Specifically, we randomly selected six samples with replacement from each of the two species and F_1_ hybrid alleles. Thus, while species identity in this approach is correct, some samples can be represented not at all or multiple times in any specific bootstrap replicate. Due to the considerable number of tested models and the associated computational burden, we restricted this bootstrapping approach to the cerebellum and cortex and performed 10 bootstrap replicates each.

### (C) Whole-brain-to-subregion comparison

We analyzed all subregions and the whole brain separately (i.e., without multivariate adaptive shrinkage) and defined a matched differential expression (DE) call as a significant DE call (FDR < 0.05) in the same direction (i.e., same sign of fold change) in the whole brain and a specific subregion. We found that 58.4 - 63.3% DE calls matched between the whole brain and any single subregion (Fig. S4A). Of the mismatched calls, 3.0 - 4.8% were mismatched because genes were either not expressed at all or filtered out due to low expression in any given subregion or the whole brain. In 10.0 - 19.5% of cases the calls did not match because there was a significant species difference in the respective subregion but not the whole brain. A possible explanation for these mismatches is overall gene expression level differences between a subregion and the whole brain that affect the power to detect DE. However, we found that the correlations of average gene expression between the whole brain and a representative subregion (here: cortex) were strong for both matched and mismatched calls (R^2^ = 0.73 vs. R^2^ = 0.69), whereas the correlation of log_2_ fold change estimates was only strong for matched (R^2^ = 0.84) but not mismatched calls (R^2^ = 0.06) (Fig. S4B). Thus, mismatches of a DE call in a subregion versus no DE in the whole brain seem not to be strongly affected by overall expression level differences between a subregion versus the whole brain. Alternatively, if these mismatched calls are mostly due to localized DE in a subregion of the brain that are not detectable (too diluted) in a whole brain sample, we would expect them to be enriched for genes that are DE in one or a few versus most or all ten subregions. Indeed, concordant with this prediction the odds of calling a gene DE in the whole brain in the same direction as a DE gene in a subregion are only 17.8% and 20.5% for a gene that is DE in a single subregion compared to 85.9% and 85.1% for a gene that is DE in all ten subregions, biased towards *P. maniculatus* or *P. polionotus*, respectively (Fig. S4C). Taken together, these results suggest that these mismatched calls (DE in subregion but not in whole brain) are mostly due to localized DE patterns. However, technical artifacts (e.g., batch effects, library preparation differences) likely contribute to these differences.

We also found 16.2 - 22.7% of cases where mismatches were due to the reverse pattern: a DE call in the whole brain but not in a given subregion. Apart from technical reasons, the most likely explanation is that these mismatches are due to DE in parts of the brain that were not captured by any of our ten subregion samples. Finally, in 0.8 - 2.2% cases were mismatches due to a “sign flip”, that is, genes were called DE in both the whole brain and a specific subregion, but the direction of expression differences (log_2_ fold changes) was inverted (Fig. S4A), which could be due to technical reasons or localized versus global (whole brain) differential expression patterns. Unfortunately, our experimental design does not permit further investigation of the cause of these mismatches.
