## Supplementary Figures and Tables for "Evolution of gene expression across brain regions in behaviorally divergent deer mice"

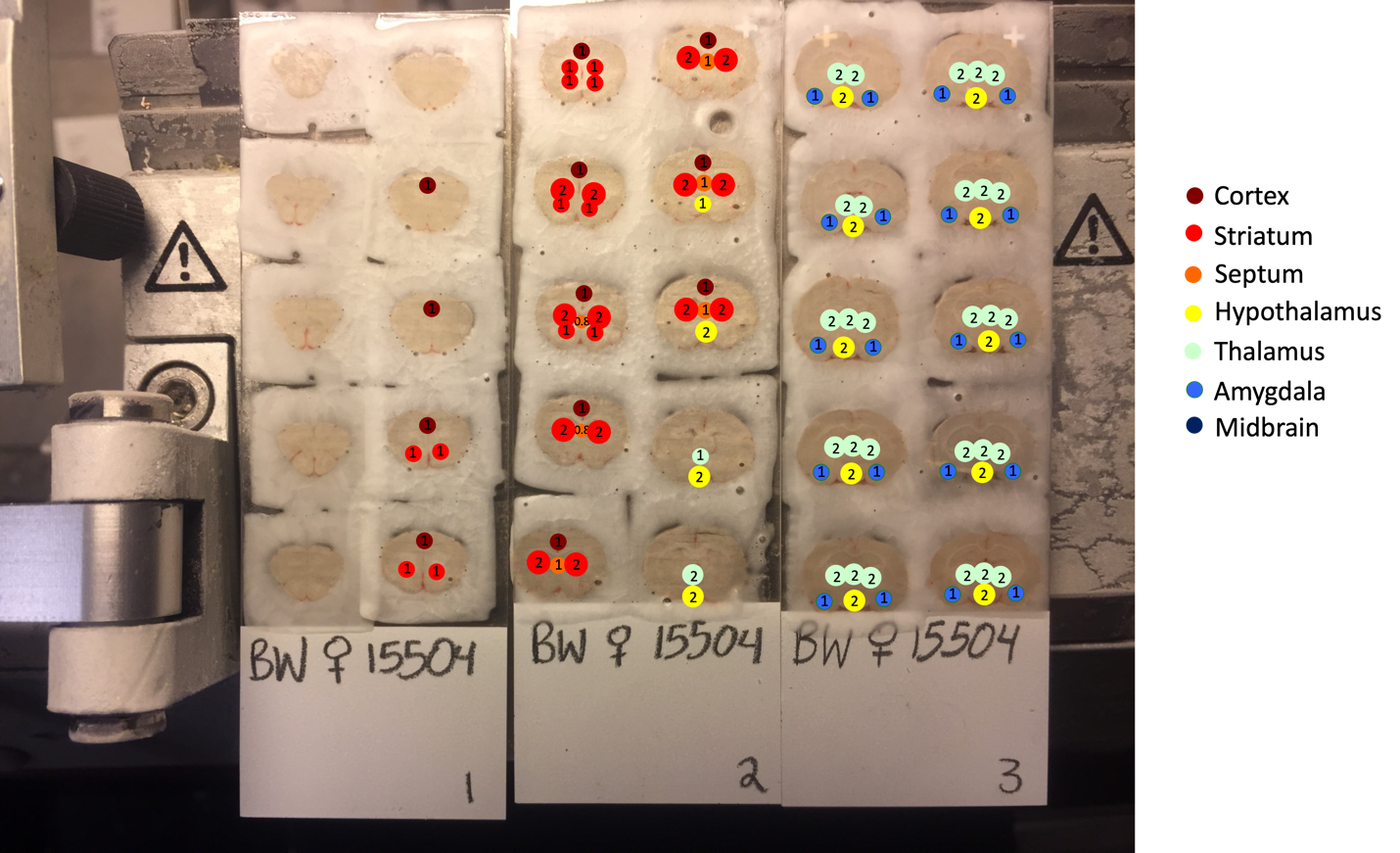

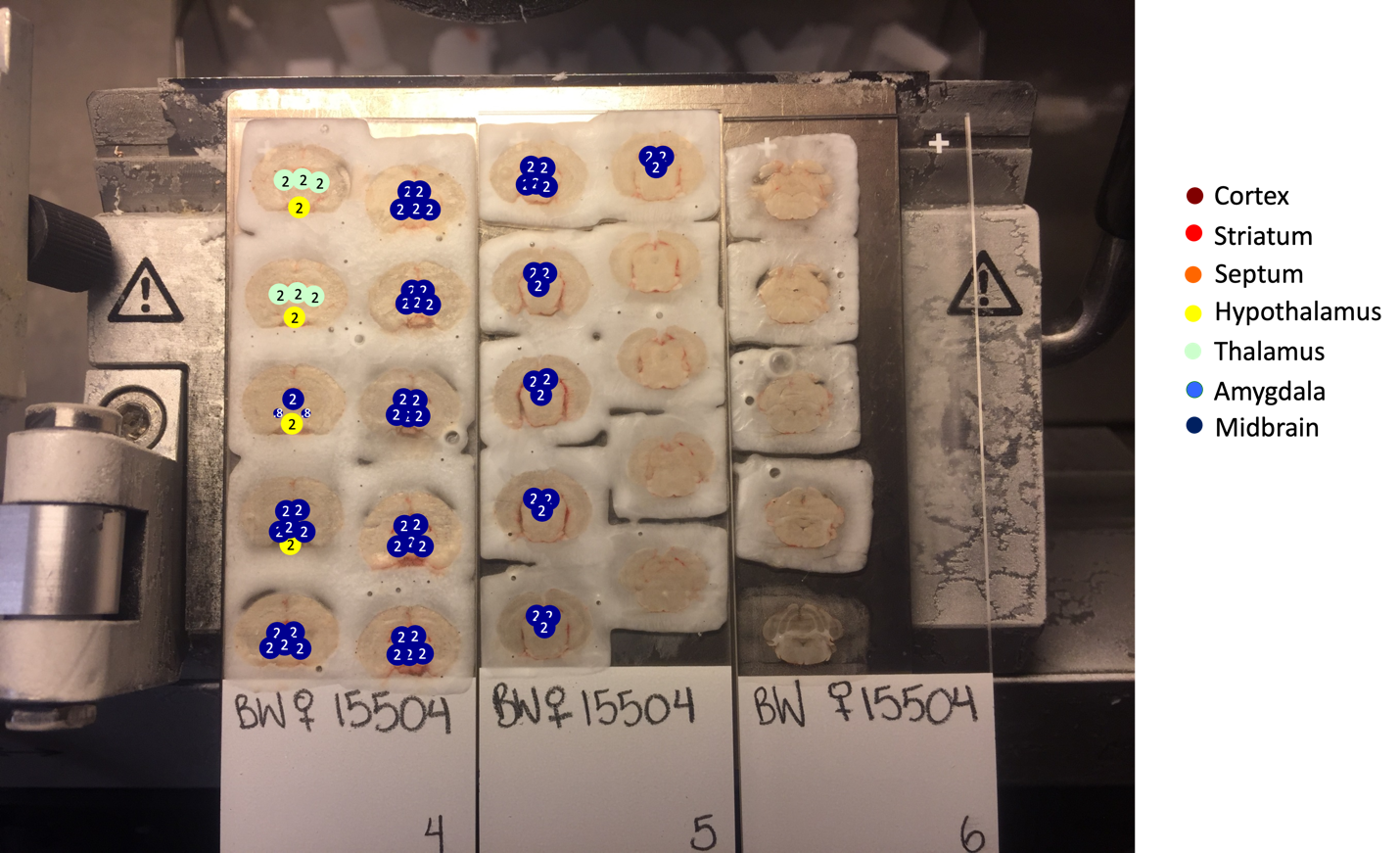

**Fig. S1. Details of subregion dissections.** Each dot represents one punch taken from a sample corer. Colors denote subregion assignment and numbers denote the punch size in mm. See Table S2 for additional details.

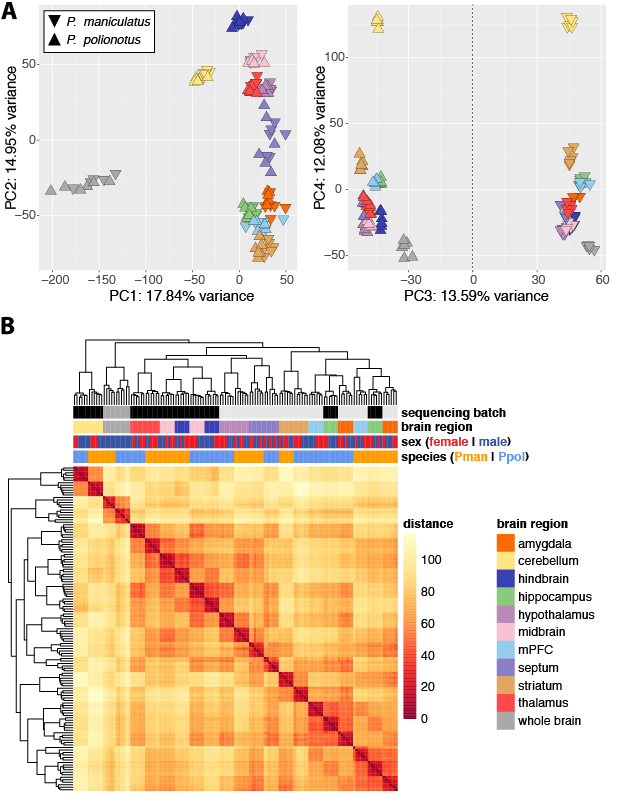

**Fig. S2. Overall variation in gene expression and sample clustering.** (**A**) Principal component analysis (PCA) of gene expression is shown PC1 x PC2 (left) and PC3 x PC4 (right). Subregions are color-coded as in panel (B) and species are indicated: *P. polionotus* (upward-pointing triangle) or *P. maniculatus* (downward-pointing triangle). Percent of variation explained by each PC is provided on axes. (**B**) Heatmap of Euclidean distance and hierarchical clustering. Sequencing batch, tissue, sex, and species are indicated for each sample. Both the PCA (A) and the clustering heatmap (B) are based on regularized log-transformed gene expression estimates.

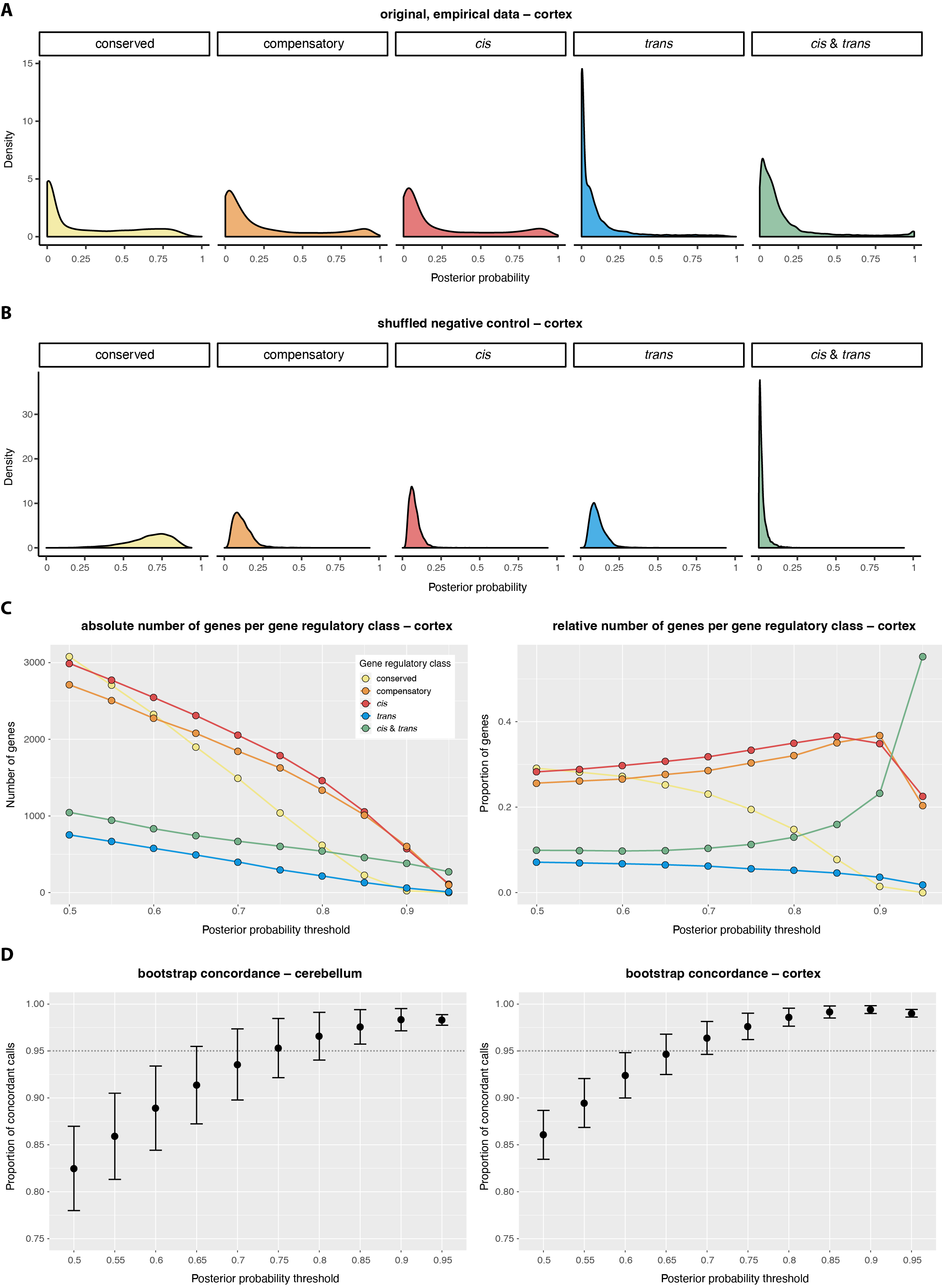

**Fig. S3. Evaluation of posterior probability thresholds.** Distributions of posterior probabilities for all five regulatory classes for the original empirical cortex data (**A**) and shuffled negative controls (**B**) in which half the samples of each species and F_1_ alleles were assigned to the other species and vice versa. (**C**) The number of genes that can be assigned to a certain gene-regulatory class with increasing posterior probability thresholds (0.5–1) in an absolute (left) and relative (right) sense. (**D**) Concordant assignments of genes to regulatory classes between the original data and ten non-parametric bootstrap replicates with increasing posterior probability thresholds (0.5–1) in the two tested subregions: cerebellum (left) and cortex (right). Highlighted is the 95% concordance level corresponding to a posterior probability threshold of 0.75.

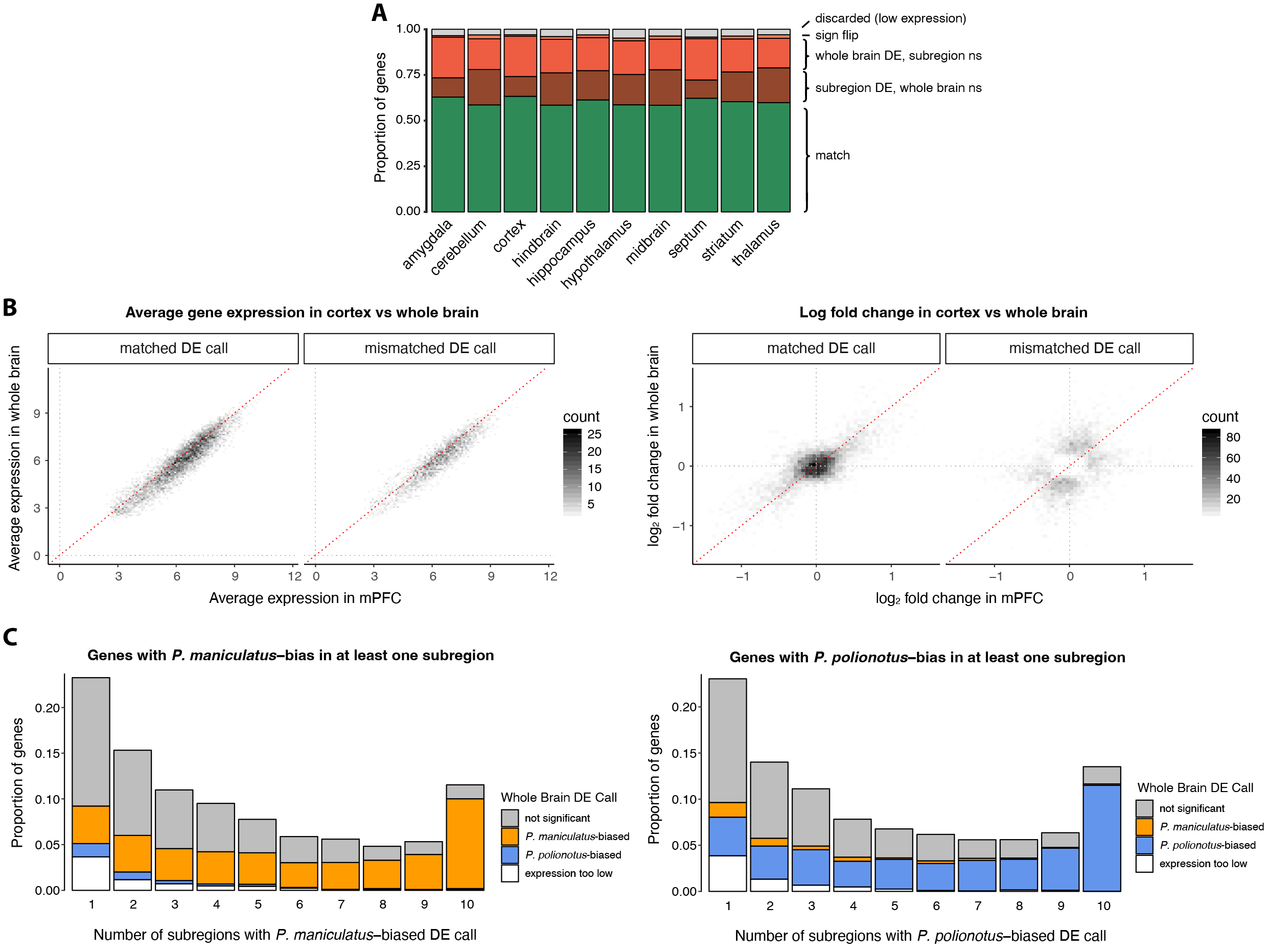

**Fig. S4. Whole-brain-to-subregion concordance.** (**A**) Summary of the proportion of genes (i) that were discarded due to low expression levels, (ii) for which a respective subregion call matched the whole brain DE call (match), (iii) that were DE in both the subregion and whole brain but with a log fold change in opposite directions (sign flip), or (iv) that differed from the whole brain calls by either being called significantly DE while the whole brain call was not significant (ns) or (v) vice versa across all subregions. (**B**) Left: average gene expression in a representative subregion (cortex) and the whole brain are highly correlated, whether their differential expression calls match or do not match. Right: in contrast, log_2_ fold change estimates are highly correlated only for genes whose DE call match but not for those that do not match. (**C**) Whole brain DE calls for genes that were either *P. maniculatus*-biased (left) or *P. polionotus-*biased (right) in at least one subregion (and not in the opposite direction) are more concordant if a gene is DE in the same direction across several subregions.

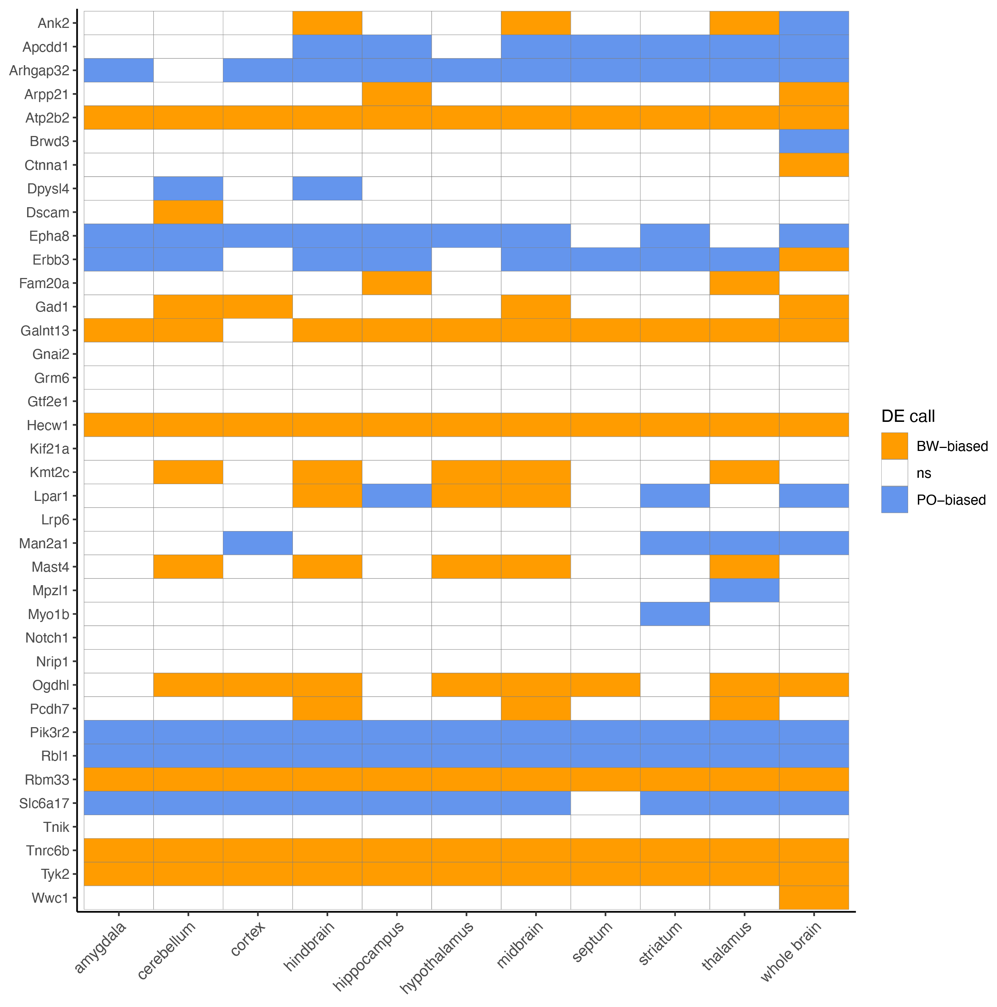

**Fig. S5. Differential expression (DE) calls of genes previously implicated in monogamy.** Each row represents one gene that had previously been suggested to be part of a universal transcriptomic mechanism underlying the evolution of monogamy in vertebrates (Young et al. 2019) and the corresponding DE call in our analyses. Note that 3 out of the 41 genes identified by Young et al. were absent in our dataset.

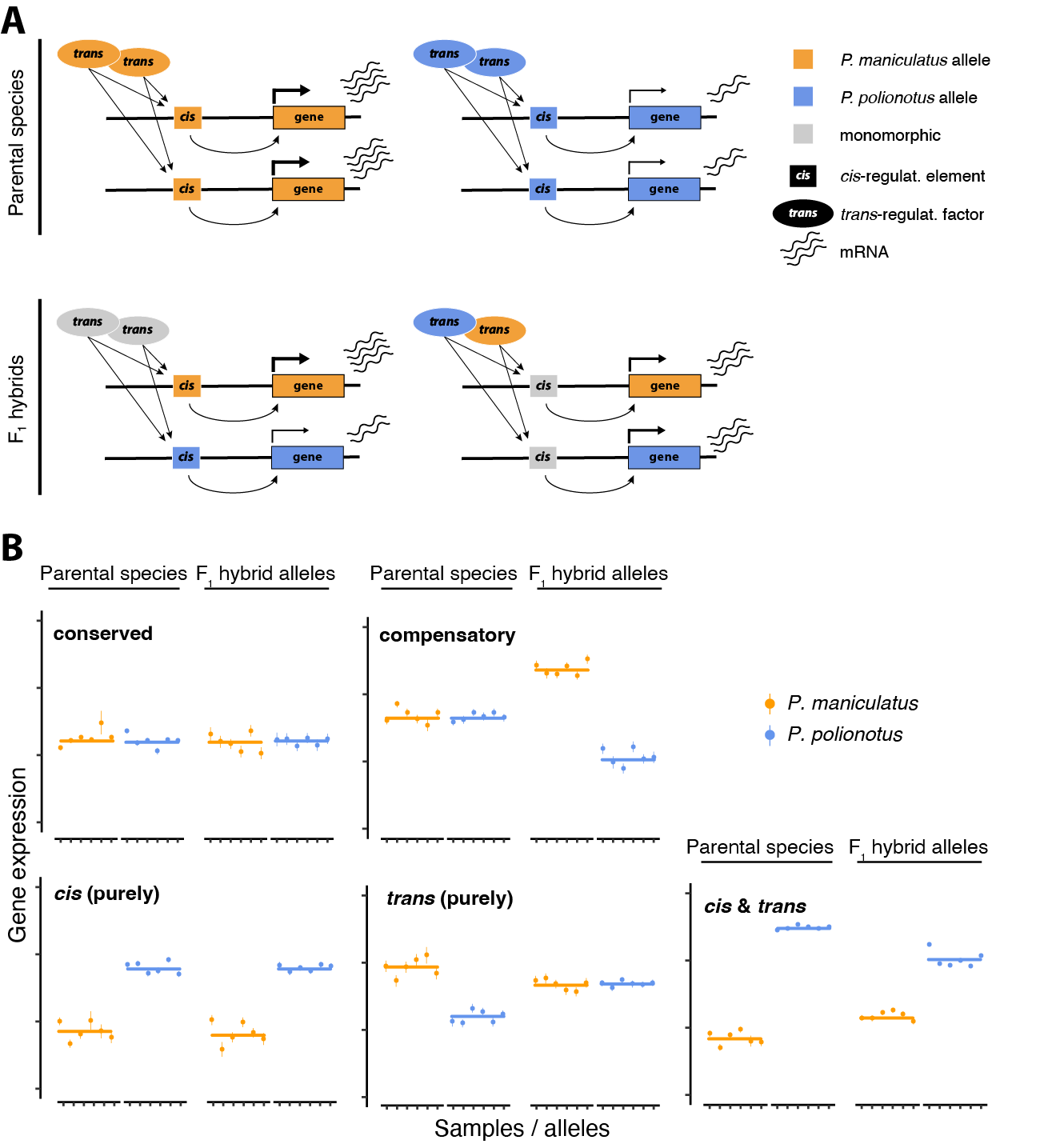

**Fig. S6. Gene regulation and allele-specific expression in F_1_ hybrids.** (**A**) The expression of genes (approximated by mRNA levels) is regulated by *cis*-regulatory elements that act in an allele-specific manner and *trans*-regulatory elements that act on both alleles. Differences in either or both elements (and there can be many for any given gene) can lead to differential gene expression between species (top). If there is no divergence of *trans*-regulatory elements, but divergence in *cis*-regulatory elements the two alleles in F_1_ hybrids may be expressed differently (bottom left). If, on the other hand, there is no divergence in *cis*-regulatory elements, but divergence in *trans*-regulatory elements, both alleles will be expressed at the same level (bottom right). (**B**) Five main regulatory classes can be distinguished by comparing the ratios of gene expression in parental species and species-specific alleles in F_1_ hybrids. If there is no differential expression between parental species, a gene’s regulation is inferred to be conserved or compensatory, depending on whether there is a difference in F_1_ hybrid alleles or not (top row). If a gene is differentially expressed between the parental species (bottom row), a gene’s regulatory divergence is inferred to be due to purely *cis*-regulatory divergence, purely *trans*-regulatory divergence, or both (*cis* & *trans*), depending on whether the expression ratio between F_1_ hybrid alleles is the same as in the parents, there is no expression difference, or there is an expression difference that deviates (is smaller or larger) from the ratio in the parental species. Illustrative gene expression estimates are shown: each dot with error bars represents a single sample’s expression estimate together with a measure of uncertainty (standard deviation of the posterior).

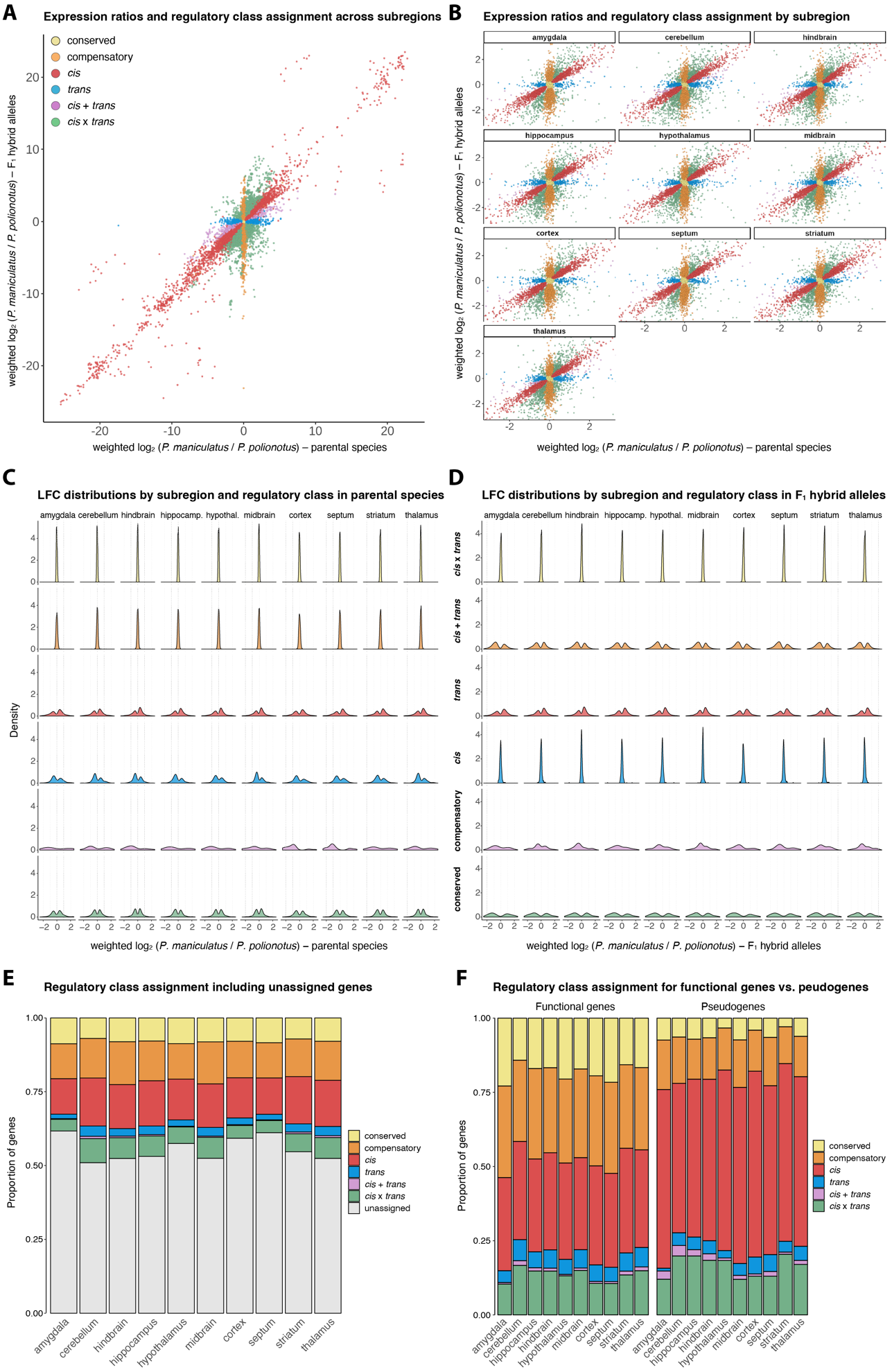

**Fig. S7. Assignment of genes to gene-regulatory classes.** (**A**) Full spectrum of weighted gene expression log_2_ fold changes in parental species versus log_2_ fold changes of alleles in F_1_ hybrids across all subregions combined, excluding pseudogenes and genes that could not be assigned to a regulatory class (N=59,091), and (**B**) subset of the spectrum for each subregion shown separately. Each dot is a gene color-coded by its inferred gene regulatory class (applying posterior probability cutoff of 0.75). (**C**) Distributions of weighted log_2_ fold change estimates for parental species and (**D**) alleles in F_1_ hybrids by gene regulatory class and subregion. (**E**) Proportion of genes assigned to different gene regulatory classes across subregions with unassigned (ambiguous) genes at the applied posterior probability cutoff. (**F**) Same as (E), but without unassigned genes and shown separately for functional genes (left) and pseudogenes (right).

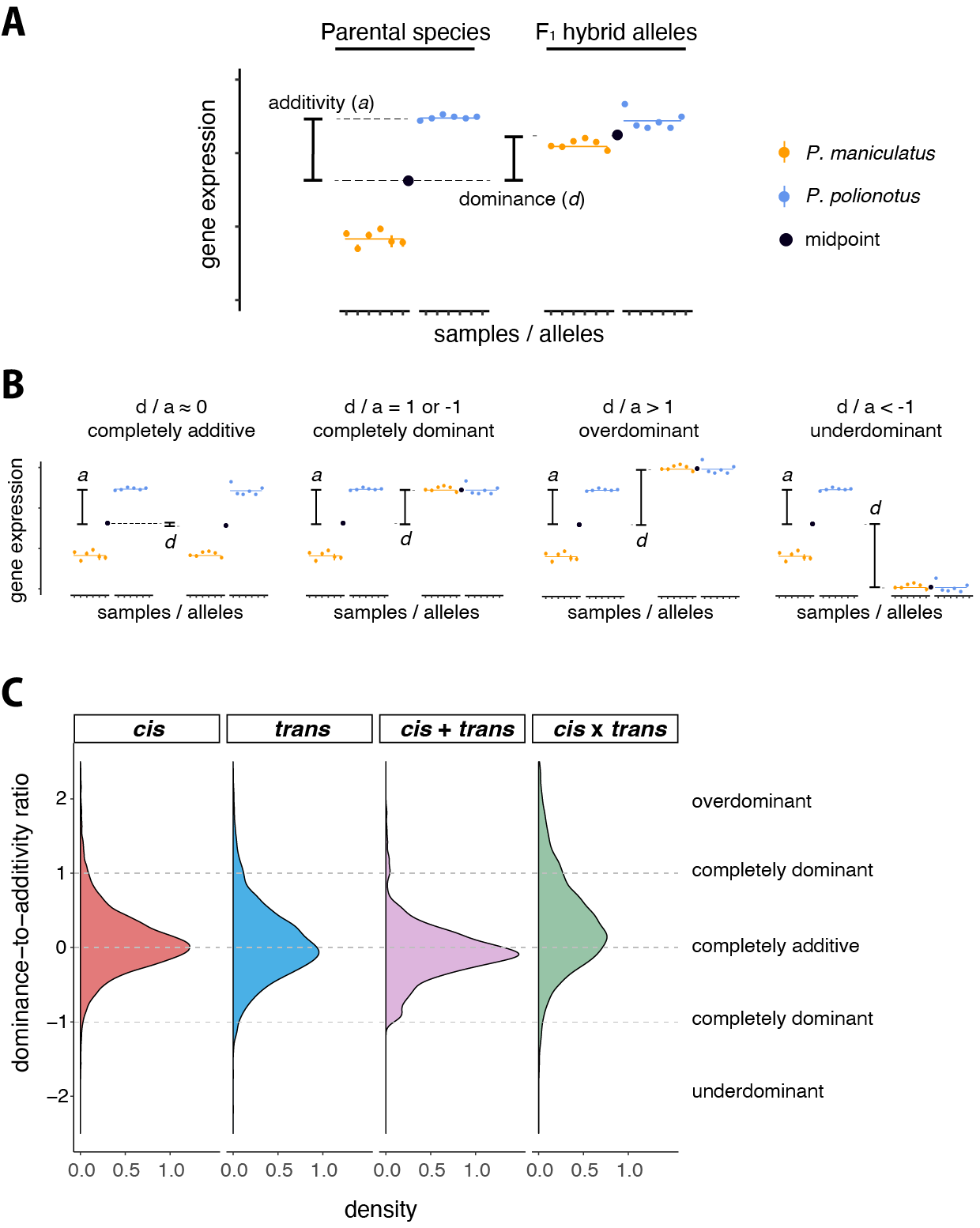

**Fig. S8. Dominance to additivity ratios.** (**A**) Example data to illustrate additivity (a) defined as half of the interparental gene expression difference and dominance (d) defined as the distance between the midpoints of parental and hybrid allele expression. (**B**) Example data to illustrate four cases: if dominance is very low compared to additivity the ratio will be close to zero and indicate complete additivity. When both *d* and *a* are approximately equal, the ratio will be close to ±1 and indicate complete dominance, in which both hybrid alleles are expressed similarly to one of the parental species. Ratios < 1 or > 1 indicate transgressive hybrid gene expression due to under- or overdominance, respectively. (**C**) Empirical distribution of dominance to additivity ratios by gene regulatory class. Ratios ~0 indicate complete additivity, ratios around ±1 complete dominance, and ratios >1 or < 1 transgressive gene expression. Note that d/a ratios for the conserved or compensatory classes were omitted, since there is no differential expression between the parental species and the additivity term thus ~0.

### **Supplementary Tables**

**Table S1. Sample overview**

| Brain region | Species / hybrid | Sex | Batch | Replicates | Sample IDs |
| --- | --- | --- | --- | --- | --- |
| cerebellum | *P. maniculatus* | female | 1 | 3 | 15508, 15526, 15690 |
| hindbrain | *P. maniculatus* | female | 1 | 3 | 15508, 15526, 15690 |
| hippocampus | *P. maniculatus* | female | 1 | 3 | 15508, 15526, 15690 |
| midbrain | *P. maniculatus* | female | 1 | 3 | 15504, 15515, 15520 |
| thalamus | *P. maniculatus* | female | 1 | 3 | 15504, 15515, 15520 |
| amygdala | *P. maniculatus* | female | 2 | 3 | 15504, 15515, 15520 |
| cortex | *P. maniculatus* | female | 2 | 3 | 15504, 15515, 15520 |
| hypothalamus | *P. maniculatus* | female | 2 | 3 | 15504, 15515, 15520 |
| septum | *P. maniculatus* | female | 2 | 3 | 15504, 15515, 15520 |
| striatum | *P. maniculatus* | female | 2 | 3 | 15504, 15515, 15520 |
| cerebellum | *P. maniculatus* | male | 1 | 3 | 15512, 15513, 15514 |
| hindbrain | *P. maniculatus* | male | 1 | 3 | 15512, 15513, 15514 |
| hippocampus | *P. maniculatus* | male | 1 | 3 | 15512, 15513, 15514 |
| midbrain | *P. maniculatus* | male | 1 | 3 | 15507, 15740, 15763 |
| thalamus | *P. maniculatus* | male | 1 | 3 | 15507, 15740, 15763 |
| amygdala | *P. maniculatus* | male | 2 | 3 | 15507, 15740, 15763 |
| cortex | *P. maniculatus* | male | 2 | 3 | 15507, 15740, 15763 |
| hypothalamus | *P. maniculatus* | male | 2 | 3 | 15507, 15740, 15763 |
| septum | *P. maniculatus* | male | 2 | 3 | 15507, 15740, 15763 |
| striatum | *P. maniculatus* | male | 2 | 3 | 15507, 15740, 15763 |
| whole brain | *P. maniculatus* | male | 3 | 5 | 16334, 16687, 16807, 16894, 16947 |
| cerebellum | *P. polionotus* | female | 1 | 2 | 15822, 15823 |
| hindbrain | *P. polionotus* | female | 1 | 2 | 15822, 15823 |
| hippocampus | *P. polionotus* | female | 1 | 2 | 15822, 15823 |
| midbrain | *P. polionotus* | female | 1 | 3 | 15764, 15818, 15821 |
| thalamus | *P. polionotus* | female | 1 | 3 | 15764, 15818, 15821 |
| amygdala | *P. polionotus* | female | 2 | 3 | 15764, 15818, 15821 |
| cortex | *P. polionotus* | female | 2 | 3 | 15764, 15818, 15821 |
| hypothalamus | *P. polionotus* | female | 2 | 3 | 15764, 15818, 15821 |
| septum | *P. polionotus* | female | 2 | 3 | 15764, 15818, 15821 |
| striatum | *P. polionotus* | female | 2 | 3 | 15764, 15818, 15821 |
| cerebellum | *P. polionotus* | male | 1 | 4 | 15765, 15819, 15820, 15920 |
| hindbrain | *P. polionotus* | male | 1 | 4 | 15765, 15819, 15820, 15920 |
| hippocampus | *P. polionotus* | male | 1 | 4 | 15765, 15819, 15820, 15920 |
| midbrain | *P. polionotus* | male | 1 | 3 | 15409, 15410, 15921 |
| thalamus | *P. polionotus* | male | 1 | 3 | 15409, 15410, 15921 |
| amygdala | *P. polionotus* | male | 2 | 3 | 15409, 15410, 15921 |
| cortex | *P. polionotus* | male | 2 | 3 | 15409, 15410, 15921 |
| hypothalamus | *P. polionotus* | male | 2 | 3 | 15409, 15410, 15921 |
| septum | *P. polionotus* | male | 2 | 3 | 15409, 15410, 15921 |
| striatum | *P. polionotus* | male | 2 | 3 | 15409, 15410, 15921 |
| whole brain | *P. polionotus* | male | 3 | 6 | 16537, 16598, 16620, 16809, 16872, 17003 |
| cerebellum | F_1_ hybrid | female | 1 | 3 | 15467, 15843, 15844 |
| hindbrain | F_1_ hybrid | female | 1 | 3 | 15467, 15843, 15844 |
| hippocampus | F_1_ hybrid | female | 1 | 3 | 15467, 15843, 15844 |
| midbrain | F_1_ hybrid | female | 1 | 3 | 15462, 15650, 15847 |
| thalamus | F_1_ hybrid | female | 1 | 3 | 15462, 15650, 15847 |
| amygdala | F_1_ hybrid | female | 2 | 3 | 15462, 15650, 15847 |
| cortex | F_1_ hybrid | female | 2 | 3 | 15462, 15650, 15847 |
| hypothalamus | F_1_ hybrid | female | 2 | 3 | 15462, 15650, 15847 |
| septum | F_1_ hybrid | female | 2 | 3 | 15462, 15650, 15847 |
| striatum | F_1_ hybrid | female | 2 | 3 | 15462, 15650, 15847 |
| cerebellum | F_1_ hybrid | male | 1 | 3 | 15845, 15846, 15848 |
| hindbrain | F_1_ hybrid | male | 1 | 3 | 15845, 15846, 15848 |
| hippocampus | F_1_ hybrid | male | 1 | 3 | 15845, 15846, 15848 |
| midbrain | F_1_ hybrid | male | 1 | 3 | 15465, 15466, 15467 |
| thalamus | F_1_ hybrid | male | 1 | 3 | 15465, 15466, 15467 |
| amygdala | F_1_ hybrid | male | 2 | 3 | 15465, 15466, 15467 |
| cortex | F_1_ hybrid | male | 2 | 3 | 15465, 15466, 15467 |
| hypothalamus | F_1_ hybrid | male | 2 | 3 | 15465, 15466, 15467 |
| septum | F_1_ hybrid | male | 2 | 3 | 15465, 15466, 15467 |
| striatum | F_1_ hybrid | male | 2 | 3 | 15465, 15466, 15467 |

**Table S2. Details for frozen dissections**

| **Order** | **Brain region** | **Slice start** | **Slice end** | **No. slices** | **No. punches** | **Punch location** | **Punch size** | **PAX^1^ start** | **PAX^1^ end** | **Tot no. slices** | **Tot no. punches** |
| --- | --- | --- | --- | --- | --- | --- | --- | --- | --- | --- | --- |
| **1** | cortex | 5/6 | 16 | 12 | 12 | medial | 1mm | 12/13 | 31/32 | 12 | 12 |
| **2** | septum | 11 | 12 | 2 | 2 | medial | 0.8mm | 21/22 | - | 6-9 | 6-9 |
| **2** | septum | 13 | 16-19 | 4 | 4 | medial | 1mm | - | 31/32 |  |  |
| **3** | striatum | 7 | 8 | 2 | 4 (2x2) | lateral | 2 x 1mm | 15/16 | - | 10 | 26 |
| **3** | striatum | 9 | 9 | 1 | 4 (2x2x1) | lateral | 2 x (1mm + 1mm) | - | - |  |  |
| **3** | striatum | 10 | 11 | 2 | 8 (2x2x2) | lateral | 2 x (2mm + 1mm) | - | - |  |  |
| **3** | striatum | 12 | 16 | 5 | 10 (5x2) | lateral | 2 x 2mm | - | 39/40 |  |  |
| **4** | thalamus | 17 | 17 | 1 | 1 | medial | 1mm | 33/34 | - | 11 | 24 |
| **4** | thalamus | 18 | 18 | 1 | 1 | medial | 2mm | - | - |  |  |
| **4** | thalamus | 19 | 20 | 2 | 4 (2x2) | lateral | 2 x 2mm | - | - |  |  |
| **4** | thalamus | 21 | 26 | 6 | 18 (3x6) | medial + lateral | 2mm + 2mm + 2mm | - | - |  |  |
| **5** | hypothalamus | 13 | 13 | 1 | 1 | medial | 1mm | 26/27 | - | 13-17 | 13-17 |
| **5** | hypothalamus | 14 | 25 | 12 | 12 | medial | 2mm | - | 49/50 |  |  |
| **6** | amygdala | 18 | 25 | 8 | 16 (2x8) | lateral | 2 x 1mm | 38/39 | 49/50 | 8 | 16 |
| **7** | midbrain | 27 | 27 | 1 | 3 | medial + lateral | 2mm + 0.8mm + 0.8mm | 52/53 | - | 11-13 | 40 |
| **7** | midbrain | 28 | 32 | 5 | 25 (5x5) | medial + lateral | 5 x (2mm) | - | - |  |  |
| **7** | midbrain | 33 | 36 | 5 | 10 (2x5) | lateral | 3 x 2mm | - | - |  |  |
| **7** | midbrain | 37 | 37 | 1 | 2 | lateral | 2 x 2mm | - | 70 |  |  |
| 1. Positions in Paxinos Mouse Brain Atlas (Paxinos & Franklin, 2004) | | | | | | | | | | | |

**Table S3. Read mapping statistics.** Given are the total number of mapped reads and the proportion of reads that mapped uniquely for each brain region sample.

| Sample ID | Brain region | Species / hybrid | total num. mapped | prop. uniquely mapped |
| --- | --- | --- | --- | --- |
| 15504 | amygdala | *P. maniculatus* | 27149105 | 0.8411 |
| 15763 | amygdala | *P. maniculatus* | 27704364 | 0.8331 |
| 15740 | amygdala | *P. maniculatus* | 28888035 | 0.8281 |
| 15515 | amygdala | *P. maniculatus* | 30616370 | 0.8291 |
| 15507 | amygdala | *P. maniculatus* | 34490220 | 0.8397 |
| 15520 | amygdala | *P. maniculatus* | 39122867 | 0.8217 |
| 15508 | cerebellum | *P. maniculatus* | 27828685 | 0.8657 |
| 15512 | cerebellum | *P. maniculatus* | 29002364 | 0.8658 |
| 15526 | cerebellum | *P. maniculatus* | 30835663 | 0.8686 |
| 15513 | cerebellum | *P. maniculatus* | 31407164 | 0.8618 |
| 15690 | cerebellum | *P. maniculatus* | 33656294 | 0.8701 |
| 15514 | cerebellum | *P. maniculatus* | 36219530 | 0.8599 |
| 15514 | hindbrain | *P. maniculatus* | 23001426 | 0.8357 |
| 15512 | hindbrain | *P. maniculatus* | 28201943 | 0.8402 |
| 15526 | hindbrain | *P. maniculatus* | 28509892 | 0.8354 |
| 15690 | hindbrain | *P. maniculatus* | 31383451 | 0.8371 |
| 15508 | hindbrain | *P. maniculatus* | 33331581 | 0.833 |
| 15513 | hindbrain | *P. maniculatus* | 33267690 | 0.8271 |
| 15513 | hippocampus | *P. maniculatus* | 26423490 | 0.829 |
| 15526 | hippocampus | *P. maniculatus* | 29125459 | 0.8203 |
| 15508 | hippocampus | *P. maniculatus* | 29403860 | 0.8229 |
| 15512 | hippocampus | *P. maniculatus* | 30258410 | 0.8325 |
| 15514 | hippocampus | *P. maniculatus* | 30731880 | 0.8302 |
| 15690 | hippocampus | *P. maniculatus* | 32959255 | 0.8376 |
| 15520 | hypothalamus | *P. maniculatus* | 29504300 | 0.824 |
| 15504 | hypothalamus | *P. maniculatus* | 30862844 | 0.8379 |
| 15515 | hypothalamus | *P. maniculatus* | 31300836 | 0.8095 |
| 15763 | hypothalamus | *P. maniculatus* | 32674503 | 0.8306 |
| 15740 | hypothalamus | *P. maniculatus* | 32984159 | 0.8284 |
| 15507 | hypothalamus | *P. maniculatus* | 37844971 | 0.8232 |
| 15504 | midbrain | *P. maniculatus* | 28933654 | 0.8398 |
| 15740 | midbrain | *P. maniculatus* | 29337334 | 0.8334 |
| 15763 | midbrain | *P. maniculatus* | 29601022 | 0.8318 |
| 15520 | midbrain | *P. maniculatus* | 32136617 | 0.8274 |
| 15515 | midbrain | *P. maniculatus* | 32915372 | 0.8371 |
| 15507 | midbrain | *P. maniculatus* | 33854619 | 0.8359 |
| 15740 | cortex | *P. maniculatus* | 12885720 | 0.6668 |
| 15763 | cortex | *P. maniculatus* | 27061045 | 0.8497 |
| 15520 | cortex | *P. maniculatus* | 28060825 | 0.8317 |
| 15515 | cortex | *P. maniculatus* | 33046334 | 0.8366 |
| 15504 | cortex | *P. maniculatus* | 35197557 | 0.8438 |
| 15507 | cortex | *P. maniculatus* | 40472898 | 0.8469 |
| 15507 | septum | *P. maniculatus* | 27762725 | 0.8308 |
| 15504 | septum | *P. maniculatus* | 28832956 | 0.8204 |
| 15763 | septum | *P. maniculatus* | 29549728 | 0.8385 |
| 15740 | septum | *P. maniculatus* | 31299257 | 0.8169 |
| 15515 | septum | *P. maniculatus* | 34300902 | 0.8187 |
| 15520 | septum | *P. maniculatus* | 50481574 | 0.8229 |
| 15520 | striatum | *P. maniculatus* | 28999371 | 0.8277 |
| 15763 | striatum | *P. maniculatus* | 29240362 | 0.8351 |
| 15515 | striatum | *P. maniculatus* | 32059680 | 0.8167 |
| 15740 | striatum | *P. maniculatus* | 33348487 | 0.814 |
| 15507 | striatum | *P. maniculatus* | 36070123 | 0.8329 |
| 15504 | striatum | *P. maniculatus* | 40976235 | 0.8326 |
| 15507 | thalamus | *P. maniculatus* | 31742533 | 0.8503 |
| 15515 | thalamus | *P. maniculatus* | 31785936 | 0.8369 |
| 15504 | thalamus | *P. maniculatus* | 32875256 | 0.8456 |
| 15763 | thalamus | *P. maniculatus* | 33303471 | 0.8495 |
| 15740 | thalamus | *P. maniculatus* | 36361156 | 0.8351 |
| 15520 | thalamus | *P. maniculatus* | 36790786 | 0.8338 |
| 16634 | whole brain | *P. maniculatus* | 17532368 | 0.8548 |
| 16947 | whole brain | *P. maniculatus* | 19672392 | 0.859 |
| 16807 | whole brain | *P. maniculatus* | 22010027 | 0.8572 |
| 16687 | whole brain | *P. maniculatus* | 24826483 | 0.8616 |
| 16894 | whole brain | *P. maniculatus* | 25668085 | 0.8454 |
| 15836 | amygdala | F1 hybrids | 55842983 | 0.5876 |
| 15847 | amygdala | F1 hybrids | 58595740 | 0.5814 |
| 15466 | amygdala | F1 hybrids | 60201290 | 0.5645 |
| 15462 | amygdala | F1 hybrids | 63454241 | 0.5912 |
| 15465 | amygdala | F1 hybrids | 66487205 | 0.5778 |
| 15650 | amygdala | F1 hybrids | 67413504 | 0.5696 |
| 15844 | cerebellum | F1 hybrids | 59870097 | 0.5765 |
| 15845 | cerebellum | F1 hybrids | 63160024 | 0.5729 |
| 15848 | cerebellum | F1 hybrids | 64659612 | 0.5789 |
| 15846 | cerebellum | F1 hybrids | 65720103 | 0.5728 |
| 15843 | cerebellum | F1 hybrids | 68702815 | 0.5721 |
| 15467 | cerebellum | F1 hybrids | 77811014 | 0.5804 |
| 15846 | hindbrain | F1 hybrids | 54442578 | 0.5371 |
| 15845 | hindbrain | F1 hybrids | 54888139 | 0.5513 |
| 15848 | hindbrain | F1 hybrids | 55969660 | 0.54 |
| 15467 | hindbrain | F1 hybrids | 61809483 | 0.543 |
| 15843 | hindbrain | F1 hybrids | 70969589 | 0.5522 |
| 15844 | hindbrain | F1 hybrids | 71722059 | 0.5327 |
| 15467 | hippocampus | F1 hybrids | 54137746 | 0.5581 |
| 15848 | hippocampus | F1 hybrids | 55344375 | 0.5658 |
| 15845 | hippocampus | F1 hybrids | 57000748 | 0.5685 |
| 15844 | hippocampus | F1 hybrids | 56951342 | 0.5636 |
| 15843 | hippocampus | F1 hybrids | 58869330 | 0.5635 |
| 15846 | hippocampus | F1 hybrids | 59381939 | 0.5444 |
| 15465 | hypothalamus | F1 hybrids | 49572042 | 0.5585 |
| 15847 | hypothalamus | F1 hybrids | 54684039 | 0.5758 |
| 15650 | hypothalamus | F1 hybrids | 55813131 | 0.5805 |
| 15836 | hypothalamus | F1 hybrids | 58832524 | 0.5729 |
| 15466 | hypothalamus | F1 hybrids | 60191298 | 0.5803 |
| 15836 | midbrain | F1 hybrids | 55629161 | 0.5483 |
| 15466 | midbrain | F1 hybrids | 62273830 | 0.5564 |
| 15465 | midbrain | F1 hybrids | 62765547 | 0.5535 |
| 15650 | midbrain | F1 hybrids | 61433150 | 0.5378 |
| 15847 | midbrain | F1 hybrids | 62629994 | 0.5395 |
| 15462 | midbrain | F1 hybrids | 65566580 | 0.5625 |
| 15465 | cortex | F1 hybrids | 47091871 | 0.559 |
| 15462 | cortex | F1 hybrids | 56051805 | 0.5835 |
| 15847 | cortex | F1 hybrids | 57109355 | 0.5722 |
| 15836 | cortex | F1 hybrids | 69739688 | 0.5696 |
| 15650 | cortex | F1 hybrids | 71915736 | 0.5542 |
| 15466 | cortex | F1 hybrids | 75355468 | 0.5799 |
| 15465 | septum | F1 hybrids | 28272361 | 0.5663 |
| 15847 | septum | F1 hybrids | 53732972 | 0.565 |
| 15466 | septum | F1 hybrids | 54685693 | 0.5638 |
| 15650 | septum | F1 hybrids | 55143270 | 0.5666 |
| 15836 | septum | F1 hybrids | 54839591 | 0.5535 |
| 15462 | septum | F1 hybrids | 62979893 | 0.5803 |
| 15650 | striatum | F1 hybrids | 59707727 | 0.5671 |
| 15836 | striatum | F1 hybrids | 64568352 | 0.5826 |
| 15847 | striatum | F1 hybrids | 84490120 | 0.5572 |
| 15465 | striatum | F1 hybrids | 96738458 | 0.555 |
| 15466 | striatum | F1 hybrids | 112042196 | 0.5764 |
| 15462 | striatum | F1 hybrids | 139812454 | 0.5766 |
| 15466 | thalamus | F1 hybrids | 52592303 | 0.5477 |
| 15836 | thalamus | F1 hybrids | 53608934 | 0.555 |
| 15847 | thalamus | F1 hybrids | 54954636 | 0.5518 |
| 15462 | thalamus | F1 hybrids | 58743673 | 0.5634 |
| 15650 | thalamus | F1 hybrids | 59824183 | 0.5461 |
| 15465 | thalamus | F1 hybrids | 64020056 | 0.5587 |
| 15409 | amygdala | *P. polionotus* | 26658857 | 0.8365 |
| 15410 | amygdala | *P. polionotus* | 29711386 | 0.8529 |
| 15821 | amygdala | *P. polionotus* | 29553711 | 0.8413 |
| 15921 | amygdala | *P. polionotus* | 30851798 | 0.8466 |
| 15818 | amygdala | *P. polionotus* | 30545278 | 0.8359 |
| 15764 | amygdala | *P. polionotus* | 33177867 | 0.8502 |
| 15822 | cerebellum | *P. polionotus* | 28480891 | 0.8698 |
| 15819 | cerebellum | *P. polionotus* | 30973108 | 0.8763 |
| 15765 | cerebellum | *P. polionotus* | 32473923 | 0.8702 |
| 15920 | cerebellum | *P. polionotus* | 34710943 | 0.868 |
| 15823 | cerebellum | *P. polionotus* | 35342148 | 0.8611 |
| 15820 | cerebellum | *P. polionotus* | 38817061 | 0.8741 |
| 15765 | hindbrain | *P. polionotus* | 25327244 | 0.8322 |
| 15819 | hindbrain | *P. polionotus* | 26400034 | 0.8252 |
| 15820 | hindbrain | *P. polionotus* | 29140677 | 0.8434 |
| 15822 | hindbrain | *P. polionotus* | 31072069 | 0.8318 |
| 15823 | hindbrain | *P. polionotus* | 32144023 | 0.8261 |
| 15920 | hindbrain | *P. polionotus* | 33267476 | 0.8413 |
| 15820 | hippocampus | *P. polionotus* | 26560684 | 0.849 |
| 15819 | hippocampus | *P. polionotus* | 26696163 | 0.852 |
| 15823 | hippocampus | *P. polionotus* | 27390193 | 0.8484 |
| 15765 | hippocampus | *P. polionotus* | 27785237 | 0.848 |
| 15822 | hippocampus | *P. polionotus* | 34308359 | 0.8538 |
| 15920 | hippocampus | *P. polionotus* | 39424735 | 0.8539 |
| 15409 | hypothalamus | *P. polionotus* | 27764939 | 0.8342 |
| 15764 | hypothalamus | *P. polionotus* | 29464364 | 0.8352 |
| 15818 | hypothalamus | *P. polionotus* | 29990600 | 0.819 |
| 15821 | hypothalamus | *P. polionotus* | 31334183 | 0.8358 |
| 15410 | hypothalamus | *P. polionotus* | 33552070 | 0.8378 |
| 15921 | hypothalamus | *P. polionotus* | 37133483 | 0.8344 |
| 15818 | midbrain | *P. polionotus* | 28904182 | 0.8366 |
| 15921 | midbrain | *P. polionotus* | 30476494 | 0.8344 |
| 15409 | midbrain | *P. polionotus* | 30874535 | 0.8366 |
| 15764 | midbrain | *P. polionotus* | 31011125 | 0.8348 |
| 15821 | midbrain | *P. polionotus* | 31729570 | 0.8344 |
| 15410 | midbrain | *P. polionotus* | 33520205 | 0.8316 |
| 15818 | cortex | *P. polionotus* | 27907729 | 0.8504 |
| 15410 | cortex | *P. polionotus* | 28114011 | 0.8315 |
| 15921 | cortex | *P. polionotus* | 30908873 | 0.8566 |
| 15821 | cortex | *P. polionotus* | 33158798 | 0.8522 |
| 15764 | cortex | *P. polionotus* | 35229535 | 0.8514 |
| 15409 | cortex | *P. polionotus* | 35715605 | 0.854 |
| 15921 | septum | *P. polionotus* | 28088999 | 0.8463 |
| 15409 | septum | *P. polionotus* | 28341283 | 0.8387 |
| 15821 | septum | *P. polionotus* | 30274308 | 0.8354 |
| 15410 | septum | *P. polionotus* | 31386137 | 0.8213 |
| 15764 | septum | *P. polionotus* | 32702954 | 0.8451 |
| 15818 | septum | *P. polionotus* | 45777503 | 0.8307 |
| 15409 | striatum | *P. polionotus* | 30353514 | 0.84 |
| 15818 | striatum | *P. polionotus* | 31501040 | 0.8298 |
| 15410 | striatum | *P. polionotus* | 31941812 | 0.836 |
| 15921 | striatum | *P. polionotus* | 32382317 | 0.8385 |
| 15821 | striatum | *P. polionotus* | 35524833 | 0.8414 |
| 15764 | striatum | *P. polionotus* | 40760359 | 0.8311 |
| 15410 | thalamus | *P. polionotus* | 30057330 | 0.852 |
| 15821 | thalamus | *P. polionotus* | 30459457 | 0.8385 |
| 15921 | thalamus | *P. polionotus* | 30757359 | 0.8455 |
| 15409 | thalamus | *P. polionotus* | 34483988 | 0.8521 |
| 15818 | thalamus | *P. polionotus* | 34992258 | 0.851 |
| 15764 | thalamus | *P. polionotus* | 75220046 | 0.8374 |
| 16598 | whole brain | *P. polionotus* | 19196127 | 0.8676 |
| 16872 | whole brain | *P. polionotus* | 20895555 | 0.8687 |
| 16620 | whole brain | *P. polionotus* | 21659133 | 0.8671 |
| 16537 | whole brain | *P. polionotus* | 25668222 | 0.8625 |
| 16809 | whole brain | *P. polionotus* | 26234923 | 0.8587 |
| 17003 | whole brain | *P. polionotus* | 28020669 | 0.8495 |
